# Redundant and cryptic enhancer activities of the Drosophila *yellow* gene

**DOI:** 10.1101/419226

**Authors:** Gizem Kalay, Jennifer Lachowiec, Ulises Rosas, Mackenzie R. Dome, Patricia Wittkopp

## Abstract

*cis*-regulatory sequences known as enhancers play a key role in regulating gene expression. Evolutionary changes in these DNA sequences contribute to phenotypic evolution. The Drosophila *yellow* gene, which is required for pigmentation, has emerged as a model system for understanding how *cis*-regulatory sequences evolve, providing some of the most detailed insights available into how activities of orthologous enhancers have diverged between species. Here, we examine the evolution of *yellow cis*-regulatory sequences on a broader scale by comparing the distribution and function of *yellow* enhancer activities throughout the 5’ intergenic and intronic sequences of *Drosophila melanogaster*, *Drosophila pseudoobscura*, and *Drosophila willistoni*. We find that *cis*-regulatory sequences driving expression in a particular tissue are not as modular as previously described, but rather have many redundant and cryptic enhancer activities distributed throughout the regions surveyed. Interestingly, cryptic enhancer activities of sequences from one species often drove patterns of expression observed in other species, suggesting that the frequent evolutionary changes in *yellow* expression observed among Drosophila species may be facilitated by gaining and losing repression of pre-existing *cis*-regulatory sequences.

## Introduction

*cis*-regulatory elements known as enhancers affect development and physiology by controlling the time, place and amount of mRNA that is transcribed from a gene. These DNA sequences range from hundreds to thousands of base pairs, are typically located in noncoding regions of the genome, and contain binding sites for transcription factors (TFs) (Spitz and Furlong 2012; Long *et al* 2016). In multicellular organisms such as Drosophila, most genes are regulated by multiple enhancers. Early studies suggested that each enhancer was responsible for a unique subset of a gene’s expression (e.g., Davidson 2001), but it is now known that multiple enhancers with varying degrees of functional redundancy can also contribute to the expression of a gene in a given cell (Perry *et al*. 2010; Frankel *et al*. 2010; Cannavò *et al*. 2016; Osterwalder *et al*. 2018; Letelier *et al*. 2018). Despite decades of research into enhancer structure and function, predicting either the location or function of enhancers from DNA sequence alone remains challenging (Lim *et al*. 2018). Consequently, empirically testing DNA sequences for their ability to activate gene expression *in vivo* using transgenic reporter genes remains a critical step for identifying enhancers and determining their function (Wittkopp and Kalay 2012; Barrière and Ruvinsky 2014).

Evolutionary changes in enhancer sequences can alter an expression pattern within a tissue, eliminate enhancer function, or create novel expression patterns, all of which can contribute to phenotypic differences within or between species (Prud’homme *et al*. 2007; Wittkopp and Kalay 2012; Rubinstein and de Souza 2013; Rebeiz and Tsiantis 2017). Modifying expression within a tissue often results from genetic changes affecting one or more transcription factor binding sites (TFBS) within an enhancer (Swanson *et al*. 2010; Rogers *et al*. 2013). Loss of enhancer activity can result from point mutations in transcription factor binding sites (Frankel *et al*. 2011) as well as larger deletions or insertions that disrupt enhancer function (Chan *et al*. 2010). It can also result from the gain of binding sites for repressors within an enhancer (Galant and Carroll 2002; Preger-Ben Noon *et al*. 2016). Genetic changes that create new enhancers are less well understood. They can evolve *de novo* from sequences that did not drive any enhancer activity previously (Eichenlaub and Ettwiller 2011; Emera *et al*. 2016) or can evolve from sequence modifications within an existing enhancer. Examples of this type of co-option have shown that sequence modifications can add activating elements (Rebeiz *et al*. 2011), eliminate repressing elements (Prabhakar *et al*. 2008; Sumiyama and Saitou 2011), or add both activating and repressing elements (Gompel *et al*. 2005; Arnoult *et al*. 2013). Cooption of existing enhancers might be more common than their *de novo* evolution because, with co-option, new activities can arise from only a small number of mutations (Gompel *et al*. 2005; Rebeiz *et al*. 2011; Arnoult *et al*. 2013; Koshikawa *et al*. 2015).

The Drosophila *yellow* gene, which encodes a protein required for the production of black pigment, has emerged as a model for studying the evolution of enhancer sequences. *yellow* expression is divergent among species, with changes in *yellow* expression evolving in concert with changes in the distribution of black melanin affecting adult body color (Wittkopp *et al*. 2002), and well-characterized enhancers controlling its expression. In *Drosophila melanogaster*, enhancers controlling *yellow* expression during the pupal stages when adult pigmentation develops have been identified in the 5’ intergenic sequence upstream of *yellow* that drive expression in the developing wings and body (head, thorax, and abdomen). An enhancer in the lone intron of *yellow* has been shown to drive *yellow* expression in bristles (Geyer and Corces 1987; Martin *et al*. 1989; Wittkopp *et al*. 2002; Jeong *et al*. 2006). Finally, a sequence required for sexually dimorphic expression in the abdomen has been identified within the body enhancer that contains binding sites for the Abdominal-B (Jeong *et al*. 2006) and Bric-a-brac (Roeske *et al*. 2018) transcription factors. Evolutionary changes in the body and wing enhancers have been identified that alter *yellow* expression in a manner that correlates with divergent pigmentation (Gompel *et al*. 2005; Jeong *et al*. 2006; Prud’homme *et al*. 2006; Kalay and Wittkopp 2010; Arnoult *et al*. 2013).

In addition to changes in individual enhancers, larger scale reorganization of enhancers within *yellow cis*-regulatory sequences have also been described, with wing and body enhancer activities found in the 5’ intergenic region of *D. melanogaster* also found in the introns of other Drosophila species (Kalay and Wittkopp 2010). This rapid evolution of the genomic organization of *yellow* enhancer activities was surprising given the collinearity of enhancers controlling expression of other genes among Drosophila species (e.g., Cande *et al*. 2009; Hare *et al*. 2008). DNA sequence analysis of *yellow cis*-regulatory regions suggested that enhancer synteny changes were due to gradual gain and loss of transcription factor binding sites rather than duplication or translocation events. Using parsimony, Kalay and Wittkopp (2010) inferred that the common ancestor of the contemporary *Drosophila* species examined most likely had wing and body enhancer activities in both the 5’ intergenic and intronic regions of *yellow*.

Here, we investigate evolutionary changes in the organization of enhancer activities within *yellow cis*-regulatory sequences by using reporter genes to test the function of sub-fragments from the 5’ intergenic and intronic sequences of *yellow* from *D. melanogaster*, *D. pseudoobscura* and *D. willistoni*. For each species, we compare enhancer activity of these sub-fragments to enhancer activity of the full region to determine which sequences are responsible for driving expression in specific parts of the fly during pupal development. We find that in all three species, redundant enhancers are common, with multiple fragments driving overlapping expression in the body and wings of developing pupae. We also find that some enhancer fragments drive cryptic expression patterns not seen in reporter genes containing larger *cis*-regulatory sequences from that species. Interestingly, these cryptic patterns are often similar to expression patterns seen in other species, suggesting that repressed enhancer activities might have contributed to the evolution of new enhancer activities. These data show that the architecture of *yellow cis*-regulatory sequences is less modular and more variable among species than previously described, with the observed organization of enhancer activities potentially making *yellow* expression more robust as well as more amenable to evolutionary change.

## Results

### Dissecting the architecture of Drosophila *yellow cis*-regulatory sequences

*D. melanogaster*, *D. pseudoobscura*, and *D. willistoni* are members of the Sophophora subgenus of Drosophila that have evolved distinct pigmentation since they diverged between 25 and 36 million years ago (Figure 1A, Russo *et al*. 1995). Specifically, *D. pseudoobscura* has evolved an overall dark pigmentation on both its abdomen and thorax, whereas *D. melanogaster* and *D. willistoni* show much more limited thoracic pigmentation and dark stripes near the posterior edge of each abdominal segment. These abdominal stripes are more lightly pigmented in *D. willistoni* than *D. melanogaster*. *D. melanogaster* males also show sexually-dimorphic dark pigment throughout the A5 and A6 abdominal segments that is absent in females, whereas neither *D. willistoni* nor *D. pseudoobscura* are considered to have sexually dimorphic pigmentation (Camino *et al*. 2015). The A6 segment of *D. melanogaster* females shows broad pigmentation in some strains (Kopp *et al*. 2003). None of these three species display the dark melanic spots of pigment found on the wings of some other *Drosophila* species (e.g., Gompel *et al*. 2005; Prud’homme *et al*. 2006; Werner *et al*. 2010; Koshikawa *et al*. 2015), but they do have dark pigment distributed evenly throughout the wing blades and wing veins. Finally, bristles throughout the body and along the wing margins are darkly pigmented in all three species. The 5’ intergenic and intronic sequences of *yellow* from all three of these species were previously shown to drive expression of a reporter gene integrated into the *D. melanogaster* genome in pupae that correlated well with these adult pigment patterns (Kalay and Wittkopp 2010).

**Figure 1.**
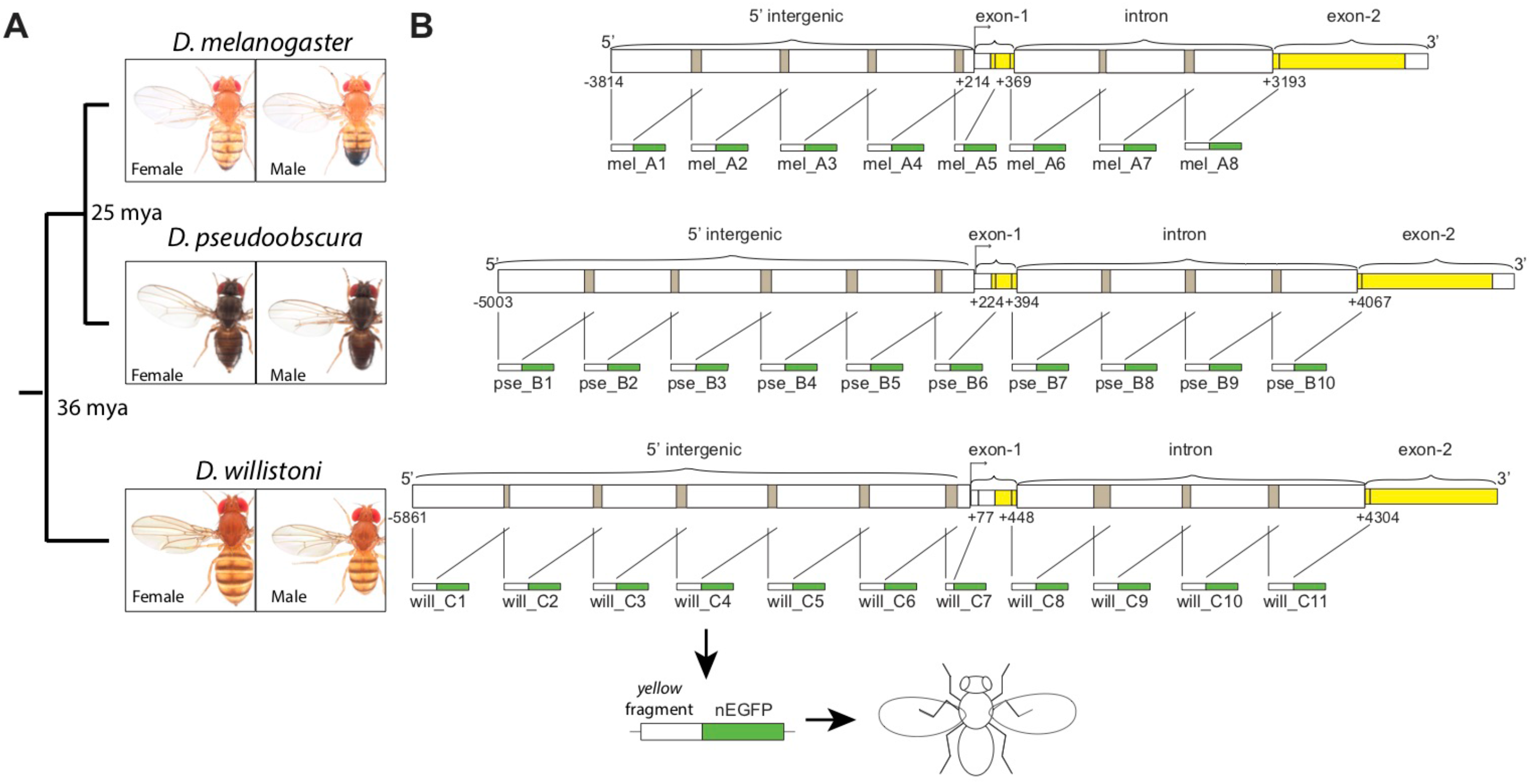
Fragments of *yellow* 5’ intergenic and intronic sequences tested for enhancer activity. (A) Divergence times among *D. melanogaster*, *D. pseudoobscura*, and *D. willistoni* are shown along with images of adult males and females from each species. (B) Schematics show the overlapping fragments from regions of 5’ intergenic and intronic sequence of *yellow* from each species tested for enhancer activity. Each of these ~1 kb fragments was cloned upstream of a nuclear Green Fluorescent Protein (nGFP) to form a reporter gene, and each reporter gene was inserted into the attP40 landing site on chromosome arm 2L. This schematic was modified from Kalay *et al*. (2016). Pictures of Drosophila species generously provided by Dr. Nicolas Gompel (Ludwig-Maximilians-University of Munich).

To determine how enhancer activities are distributed within the previously tested 5’ intergenic and intronic regions of *yellow*, we subdivided each full region into ~1 kb fragments with adjacent fragments overlapping by about 100 bp (Figure 1B, Supplementary File 1). Each of these fragments was cloned upstream of a minimal hsp70 promoter and the coding sequence for a nuclear Enhanced Green Fluorescent Protein (nEGFP) and transformed into the same site of the *D. melanogaster* genome using the attB/attP system for targeted insertion (Bischof *et al*. 2007; Markstein *et al*. 2008). Enhancer activity was assessed using confocal microscopy in flies ~80 hours after puparium formation. We analyzed this stage because it is when *yellow* expression is required for development of adult pigmentation (Massey and Wittkopp 2016); additional expression might be driven by these enhancers at other developmental stages.

Using these confocal images of pupal bodies, we characterized enhancer activity in the epidermal and bristle cells of the head, thorax and abdomen, recording whether or not the expression pattern among replicate flies consistently showed (1) stripes of expression in abdominal segments, (2) broad expression within abdominal segments, (3) sexually dimorphic expression in abdominal segments A5 and A6, (4) expression marking the circumference of the head, (5) expression on the top of the head, (6) expression in the thoracic trident pattern, (7) expression in the thoracic flight muscle attachment sites (Fernandes *et al*. 1996), and/or (8) expression in bristles on the body (Figure 2A). In the wings, we recorded whether there was expression in the (1) epidermal cells of the wing blade, (2) longitudinal and cross veins, and/or (3) bristles along the anterior margin (Figure 2A). We inferred enhancer activity when we consistently saw higher expression driven by a reporter gene in a tissue than that driven by a control reporter gene lacking any *yellow* fragment (Figure 2B,C). Fluorescence in the control line was highest and most variable in the wing blade (Supplementary Figure 1), making it difficult to confidentially infer enhancer activities that drive low expression in this part of the fly. In all cases where expression elements were difficult to interpret, we report the opinion of the majority of co-authors. All transformant flies showed high expression levels of Green Fluorescent Protein (GFP) in the eyes and ocelli driven by the *Pax6-GFP* transformation marker (Figure 2, Horn and Wimmer 2000).

**Figure 2.**
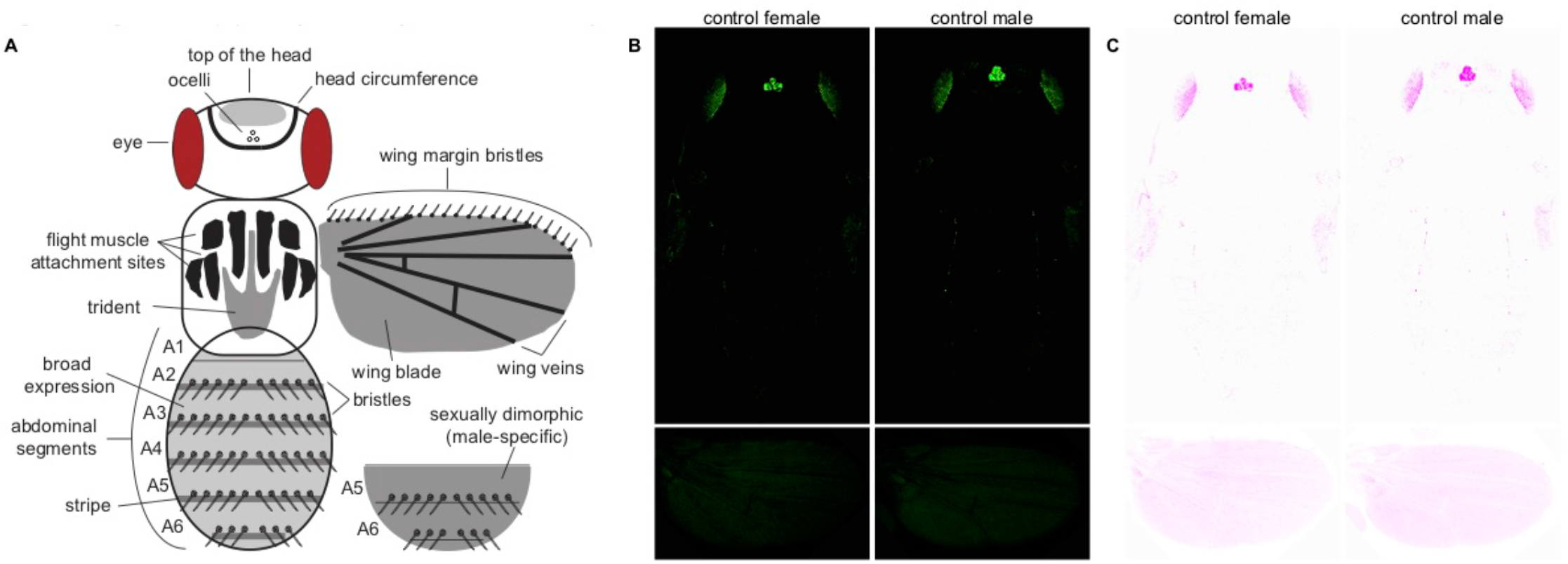
Regions of the pupal body scored for *yellow* enhancer activity. (A) This schematic represents a Drosophila pupa and has the regions scored for reporter gene expression on the head, thorax, abdomen, and wings indicated. Abdominal segments A1 – A6 are also labeled. Eyes and ocelli are also shown because the transformation marker (Pax6-GFP) inserted with the *yellow* fragment reporter genes drives expression in these tissues. (B) Activity of the reporter gene without any *yellow* enhancer fragment. These images show background activity of the “empty” reporter gene. The fluorescence observed in the eyes and ocelli is driven by the Pax6-GFP transformation marker that was integrated with the empty reporter gene. (C) The same control flies as in (B) are shown here using an inverted color scheme. This inverted color scheme makes it easier to see low levels of fluorescence.

For 30 of the 35 transformant lines, we imaged individuals homozygous for the transgene. For the five remaining lines (mel_A5, pse_B4, pse_B6, pse_B7, and will_C8, indicated with asterisks in Figures 4-6), we were unable to recover homozygous flies and imaged heterozygous individuals instead. These heterozygous flies showed weaker expression of GFP in the eyes and ocelli, suggesting that our images for these lines underestimated expression relative to homozygous individuals in other tissues. Some variability in fluorescence levels among images was also expected to result from fluctuations in confocal laser intensity despite the fact that we used the same laser and imaging settings each day. Expression of the green fluorescent reporter gene is shown using an inverted color scheme in Figures 3-5 to make it easier to see low levels of expression. The fluorescent images, which show differences in regions of high expression more clearly, are also provided as Supplementary Figures 2-4.

### Pupal enhancer activities of *D. melanogaster yellow*

The 4 kb intergenic sequence located 5’ of the *D. melanogaster yellow* gene drove expression in the thoracic trident and abdominal stripes (Figure 3A, Kalay and Wittkopp 2010). Sexually dimorphic expression was also observed in the abdomen, with expression expanded and elevated in the A5 and A6 segments of males relative to females. Females also showed broader expression in the A6 segment than segments A1-A5. Among the subfragments from the 5’ intergenic sequence tested (Figure 3A), the mel_A2 fragment drove stripes of expression in the abdomen most similar to the full 5’ intergenic sequence, but broad expression was limited only to the A6 segment of males. Sexually dimorphic expression in abdominal segments A5 and A6 was most similar to that observed with the full enhancer for fragments mel_A1 and mel_A4, but these fragments also drove broad expression throughout segments A2-A4. The mel_A3 fragment drove high levels of expression throughout the abdomen, with the highest levels seen throughout segments A5 and A6 and in a stripe near the posterior edge of segments A2-A4 in both males and females. In the thorax, a trident pattern of expression similar to that driven by the full 5’ intergenic sequence was observed for mel_A1, mel_A2, and mel_A4. For mel_A3, strong expression was driven throughout the thorax (including thoracic trident and flight muscle attachment sites). In the head, expression in the circumference was most prominent for the mel_A2 and mel_A3 fragments, with lower levels of expression also observed on the top of the head for mel_A2, mel_A3, and mel_A4 (Figure 3A). In the wings (Figure 3B), the full 5’ intergenic sequence drove expression in the epidermal cells of the wing blade. Expression in the wing blade was also driven by mel_A1, mel_A2, and mel_A3, with the highest expression driven by mel_A2. Expression in the wing blade was consistently elevated in the region surrounding and posterior to the L5 wing vein for the mel_A1 fragment (arrows, Figure 3B). Finally, one fragment, mel_A4 also drove expression at low levels in the wing veins. Fragment mel_A4 might also drive low levels of expression in the wing (Figure 3B), but this enhancer activity was not called because some control flies showed similar levels of fluorescence (Supplementary Figure 1). In all of the tissues examined, the mel_A5 fragment, which was the shortest, assessed in flies heterozygous for the transgene, and closest to the transcription start site of *yellow*, failed to drive expression of the reporter gene above background levels.

**Figure 3.**
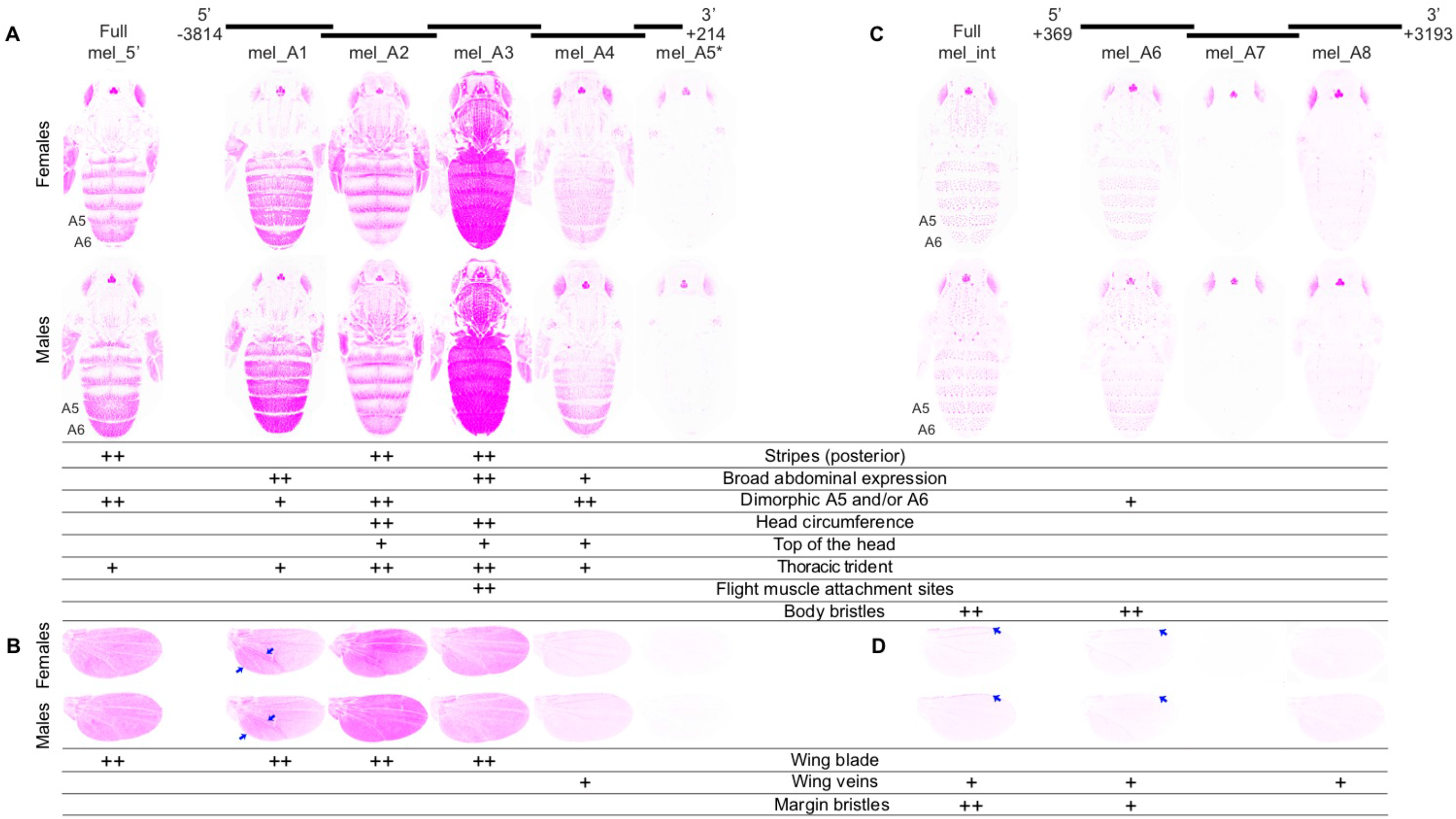
Enhancer activity of *D. melanogaster yellow* fragments. Below the schematic of overlapping sequence regions from the 5’ intergenic and intronic regions of the *D. melanogaster yellow* gene tested for enhancer activity are images of transgenic pupae that show expression from the GFP reporter genes in magenta. Expression driven by fragments from the 5’ intergenic region (A,B) and the intron (C,D) is shown in the body (A,C) and wings (B,D) for both female (top row) and male (bottom row) flies. Summary tables below each pair of pupal bodies (A,C) and wings (B,D) show our interpretation of these images: “++” indicates strong fluorescence observed in the body region, whereas “+” indicates weaker fluorescence. Abdominal segments A5 and A6 are indicated on the body images showing expression driven by the full 5’ intergenic and intronic sequences. Blue arrows highlight elevated expression in a region of the wing driven by the mel_A1 fragment and expression in the margin bristles driven by the full *D. melanogaster* intron (“Full mel_int”) and the mel_A6 fragment. The asterisk next to the mel_A5 fragment indicates that activity of this element is shown for flies heterozygous for the reporter gene. All other images show GFP expression in flies homozygous for the reporter gene. The magenta color used in this figure makes it easier to see low expression levels; a copy of this figure with GFP expression shown in the more traditional green is provided as Supplementary Figure 2.

The 2.7 kb intronic sequence of the *D. melanogaster yellow* gene drove expression in the bristles of the body, but not epidermal cells (Figure 3C, Kalay and Wittkopp 2010). Among the intronic fragments (Figure 3C), sequences included in the mel_A6 fragment were sufficient to drive expression in this pattern, although expression seemed to be lower (especially in A2 and A3). The mel_A6 fragment also appeared to drive low levels of sexually dimorphic expression in A5 and A6 in males. Neither the mel_A7 nor mel_A8 fragments drove expression in any of the body regions scored. In the wings (Figure 3D), the full intronic sequence drove low levels of expression in the veins, with higher levels of expression in the margin bristles (arrows, Figure 3D). Fragments mel_A6 and mel_A8 also drove low levels of expression in the wing veins, with mel_A6 driving expression in the margin bristles as well (arrows, Figure 3D). The other intronic fragment, mel_A7, did not drive any detectable expression in the wing.

### Pupal enhancer activities of *D. pseudoobscura yellow*

The 5.3 kb intergenic sequence located 5’ of the *D. pseudoobscura yellow* gene drove expression throughout the developing head, thorax, and abdomen (Figure 4A, Kalay and Wittkopp 2010). Although pigmentation of this species is generally considered to be sexually monomorphic (Kopp *et al*. 2000; Jeong *et al*. 2006; Salomone *et al*. 2013; Camino *et al*. 2015), expression was elevated in the A5 and A6 abdominal segments relative to A2-A4 in males and decreased in A6 relative to A2-A5 in females. Among the sub-fragments from the 5’ intergenic sequence tested (Figure 4A), pse_B1, pse_B3, and pse_B5 all drove broad expression in the body similar to the full 5’ intergenic region; however, of these three fragments, only pse_B1 drove sexually dimorphic expression similar to the full 5’ intergenic sequence. Sexually dimorphic expression in abdominal segments A5 and A6 was also driven by pse_B2, but without the accompanying broader abdominal expression. This fragment also drove expression at the lateral edges of segment A4 in males and at the posterior half of segments A5 and A6 in females. The pse_B3 fragment also drove elevated expression along the posterior edge of each abdominal segment near the midline in females (arrows, Figure 4A), but we did not score this expression as “stripes” because it was not throughout the segment nor present in males. In addition, the pse_B3 fragment drove a lower level of expression in the A6 abdominal segments of both males and females; expression was also reduced to a lesser extent in the A5 abdominal segment of males. In the thorax, the full 5’ intergenic sequence drove low levels of expression in the trident as well as higher levels of expression along the lateral edges of the thorax. Among the intergenic sub-fragments, the pse_B3 fragment drove expression in the most similar pattern, albeit with lower expression in the middle of the thorax and with expression also in flight muscle attachment sites. The pse_B1 fragment also consistently drove expression along the lateral edges of the thorax, but expression in the trident pattern was much more variable among replicate flies. Stronger thoracic expression was driven by pse_B4 and pse_B5 in both the trident and flight muscle attachment sites. Although the only detectable expression driven by pse_B4 was in the thorax, it might also drive low levels of expression in the abdomen that were missed by assessing activity of this fragment only in a heterozygous state. In the head, pse_B5 drove expression in the head circumference, whereas pse_B1 and pse_B3 both drove expression on the top of the head more similar to the full 5’ intergenic sequence. In the wings (Figure 4B), the full 5’ intergenic fragment drove low levels of epidermal cell expression throughout the wing blade without any expression in the wing veins or margin bristles. Similar epidermal expression throughout the wing blade was driven by pse_B1, pse_B2, pse_B3, pse_B4, and pse_B5. All of these fragments except pse_B4 also drove ectopic expression in the veins. The pse_B6 fragment, which was the shortest, assessed in flies heterozygous for the transgene, and closest to the transcription start site of *yellow*, did not drive expression in any of the pupal structures scored (Figure 4A,B).

**Figure 4.**
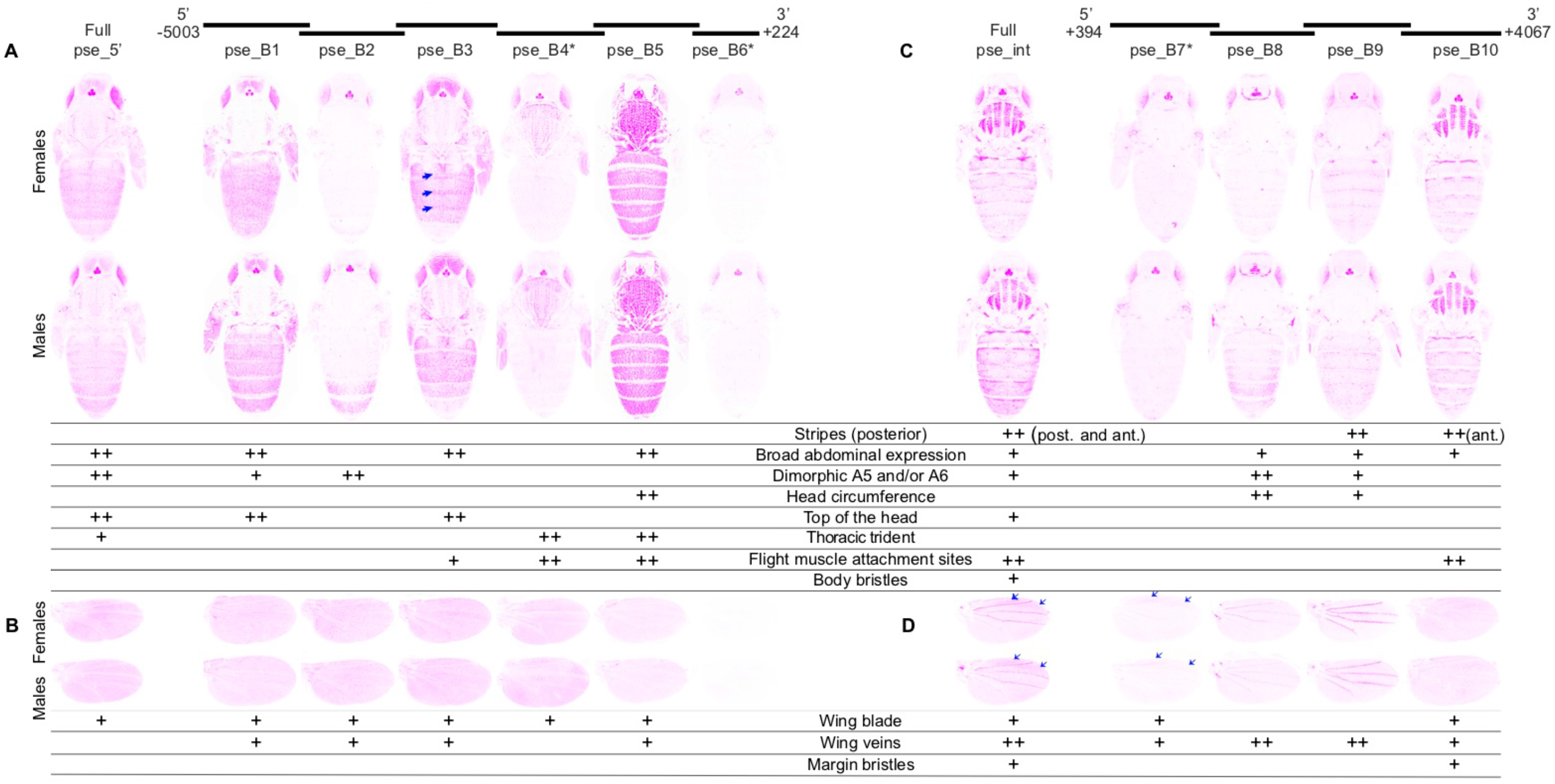
Enhancer activity of *D. pseudoobscura yellow* fragments. Below the schematic of overlapping sequence regions from the 5’ intergenic and intronic regions of the *D. pseudoobscura yellow* gene tested for enhancer activity are images of transgenic pupae that show expression from the GFP reporter genes in magenta. Expression driven by fragments from the 5’ intergenic region (A,B) and the intron (C,D) is shown in the body (A,C) and wings (B,D) for both female (top row) and male (bottom row) flies. Summary tables below each pair of pupal bodies (A,C) and wings (B,D) show our interpretation of these images: “++” indicates strong fluorescence observed in the body region, whereas “+” indicates weaker fluorescence. Note that in addition to the stripes seen at the posterior edge of abdominal segments in many cases, the full *D. pseudoobscura* intron (“Full pse_int”) as well as fragment pse_B10 drove elevated expression in stripes at the anterior edge of each abdominal segment indicated by the “ant.” notation in the table. Blue arrows highlight elevated expression along the posterior edge of abdominal segments in females driven by the pse_B3 fragment and between the L1 and L2 as well as L2 and L3 veins in the wing blade driven by full pse_int and the pse_B7 fragment. The asterisk next to the pse_B4, pse_B6, and pse_B7 fragments indicates that activity of this element is shown for flies heterozygous for the reporter gene. All other images show GFP expression in flies homozygous for the reporter gene. The magenta color used in this figure makes it easier to see low expression levels; a copy of this figure with GFP expression shown in the more traditional green is provided as Supplementary Figure 3.

The 3.6 kb intronic sequence of the *D. pseudoobscura yellow* gene drove expression in the flight muscle attachment sites of the thorax, on top of the head, and throughout each abdominal segment, with expression elevated at the anterior and posterior edges of each abdominal segment (Figure 4C, Kalay and Wittkopp 2010). Broad, male-specific expression in abdominal segments A5 and A6 as well as along the lateral edges of segments A2-A4 was also observed. Expression in bristles on the body was driven by the full intron too, but this expression was much more subtle than the bristle expression driven by the *D. melanogaster* intron (Figure 3C). Among the intronic sub-fragments (Figure 4C), pse_B8, pse_B9, and pse_B10 all drove subsets of the abdominal expression pattern driven by the full intron. Specifically, pse_B8 drove broad expression elevated along the lateral edges of A2-A4 and throughout segments A5 and A6 of males. The pse_B9 fragment drove this sexually dimorphic expression pattern as well, but more weakly, and also drove elevated expression at the posterior edge of each abdominal segment. The pse_B10 fragment drove expression throughout the abdomen, with expression elevated in stripes along the anterior edge of each abdominal segment. None of the intronic fragments drove expression in the body bristles similar to that driven by the full intron. By contrast, the thoracic expression driven by the full intron was mimicked by pse_B10, with strong expression in the flight muscle attachment sites. Expression on the top of the head driven by the full intron was not consistently reproduced by any of the sub-fragments; however, fragments pse_B8 and pse_B9 drove strong expression in the head circumference. Low levels of expression driven by pse_B7 may have been missed because this reporter gene was heterozygous in the flies assayed. In the wings, the full intronic fragment drove expression in all of the structures scored (blade, veins, and bristles), with elevated expression in the wing blade between the L1 and L2 as well as L2 and L3 veins (arrows, Figure 4D). As in the body, expression in bristles along the wing margin was more subtle than that driven by the *D. melanogaster* intron. Among the sub-fragments (Figure 4D), pse_B7 drove expression between the L1 and L2 as well as L2 and L3 veins similar to the regions of highest expression driven by the full intron (arrows, Figure 4D), and pse_B10 drove low levels of expression in the wing blade (Figure 4D). Expression in the veins was driven by all four intronic sub-fragments, with the highest expression driven by the pse_B8 and pse_B9 fragments. Finally, expression in the wing margin bristles was driven by the pse_B10 fragment.

### Pupal enhancer activities of *D. willistoni yellow*

The 5.9 kb *D. willistoni yellow* 5’ intergenic region drove expression in the abdomen highest in stripes along the posterior edge of segments A2-A5 in both males and females with expression reduced in A5 and undetectable in A6 (Figure 5A, Kalay and Wittkopp 2010). Among the sub-fragments from the 5’ intergenic sequence (Figure 5A), will_C1 and will_C3 drove stripes of expression elevated at the posterior edge of each segment. The will_C4 fragment also drove elevated expression in stripes at the posterior edge of most segments, with some elevated expression also seen near the anterior edges of some segments. Broad expression throughout abdomen segments was driven by will_C1, will_C2, and will_C4 as well as (to a lesser extent) will_C6 and will_C7. Surprisingly, sexually dimorphic patterns of elevated expression in the A5 and/or A6 of males relative to females were driven by will_C1, will_C2, will_C3, and will_C4 despite the absence of sexually dimorphic pigmentation or expression driven by the full 5’ intergenic sequence. In the thorax, the full 5’ intergenic sequence drove expression in the trident as well as the surrounding flight muscle attachment sites. Similar patterns of expression were driven at higher levels by the will_C1 and will_C3 fragments, with will_C4 driving lower expression detected only in the trident. No expression in the head appeared to be driven by the full 5’ intergenic fragment, yet will_C2 drove strong expression on the top of the head, will_C4 drove some low expression in the head circumference, and will_C1 drove expression in both of these areas. In the wings (Figure 5B), the full 5’ intergenic sequence drove expression most clearly in the wing veins. This sequence might also drive low expression throughout the wing blade (Figure 5B), but we did not call this enhancer activity because some images from the control line had similar fluorescence (Supplementary Figure 1). All of the sub-fragments except will_C5 drove expression in the wing-blade above background levels, with the highest expression levels driven by will_C1, will_C3, and will_C6. Expression in the wing blade driven by the will_C2 fragment was highest in proximal areas of the wing (arrows, Figure 5B). The highest levels of expression in the veins were driven by the will_C5 fragment, but expression in veins was also observed for fragments will_C4 and will_C7.

**Figure 5.**
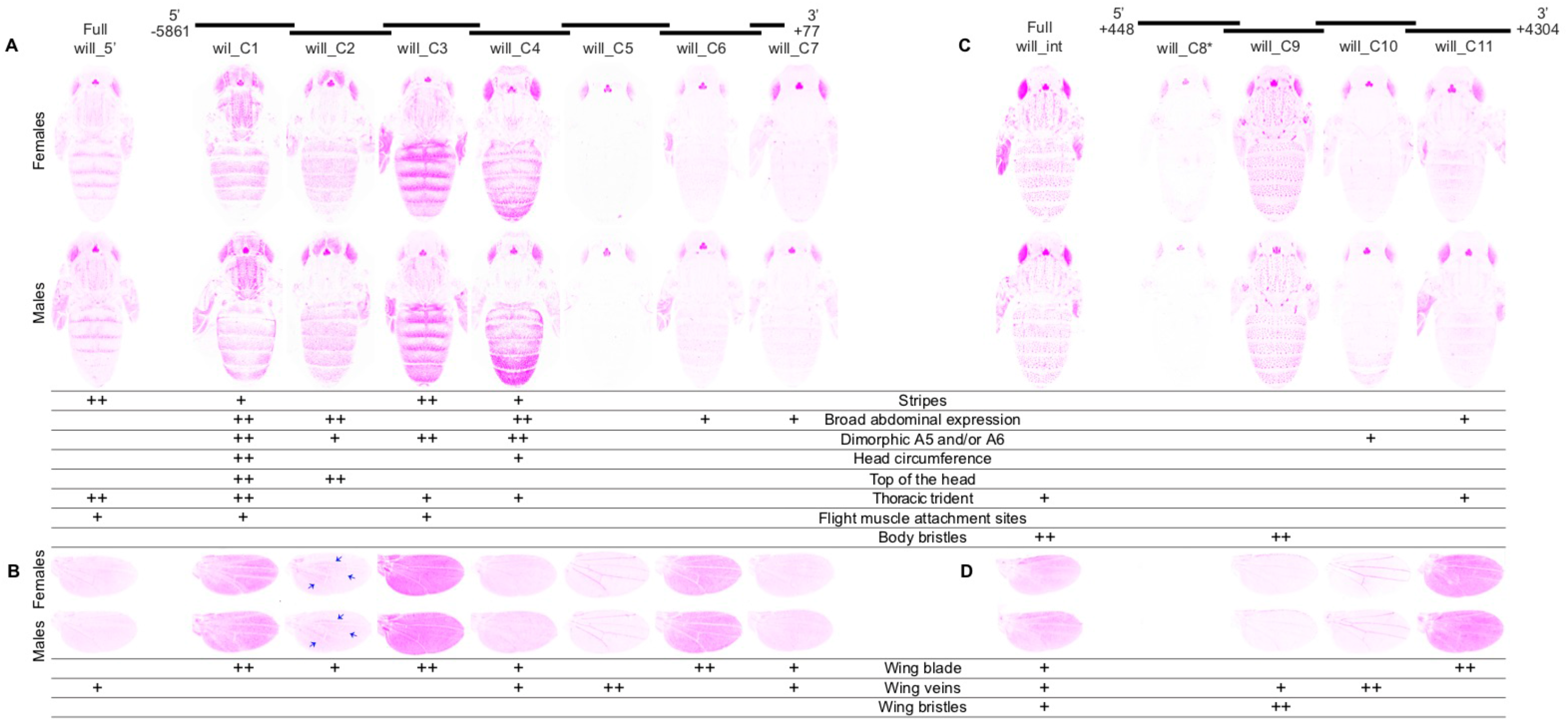
Enhancer activity of *D. willistoni yellow* fragments. Below the schematic of overlapping sequence regions from the 5’ intergenic and intronic regions of the *D. willistoni yellow* gene tested for enhancer activity are images of transgenic pupae that show expression from the GFP reporter genes in magenta. Expression driven by fragments from the 5’ intergenic region (A,B) and the intron (C,D) is shown in the body (A,C) and wings (B,D) for both female (top row) and male (bottom row) flies. Summary tables below each pair of pupal bodies (A,C) and wings (B,D) show our interpretation of these images: “++” indicates strong fluorescence observed in the body region, whereas “+” indicates weaker fluorescence. Blue arrows highlight elevated expression in the proximal areas of the wing driven by the will_C2 fragment. The asterisk next to the will_C8 fragment indicates that activity of this element is shown for flies heterozygous for the reporter gene. All other images show GFP expression in flies homozygous for the reporter gene. The magenta color used in this figure makes it easier to see low expression levels; a copy of this figure with GFP expression shown in the more traditional green is provided as Supplementary Figure 4.

The 3.7 kb *D. willistoni yellow* intronic region drove expression most prominently in the bristle cells of the body and epidermal cells in the thoracic trident (Figure 5C, Kalay and Wittkopp 2010). Among the sub-fragments (Figure 5C), will_C9 fully recapitulated the bristle expression whereas will_C11 drove expression in the thoracic trident. Low levels of broad expression in the abdominal segments were driven by the fragment will_C11 despite no clear epidermal expression driven by the full intron (Figure 5C). The will_C10 fragment drove sexually dimorphic expression in abdominal segments A5, A6 and the lateral edges of A4 that was not clearly seen in expression driven by the full intron. In the wings, the full intronic fragment drove expression throughout the wing blade, in the margin bristles, and at low levels in the veins (Figure 5D). Among the sub-fragments (Figure 5D), expression in the wing blade was driven solely by fragment will_C11, and margin bristle expression was driven solely by fragment will_C9. Expression in the veins was driven by fragments will_C9 and will_C10. The will_C8 fragment, whose enhancer activity was assessed in heterozygous flies, did not drive detectable expression in any of the body or wing structures scored.

### Independent elements of abdominal expression patterns identified by principal component analysis

To complement our qualitative assessments of *yellow* enhancer activity, we also used morphometric tools combined with principal component analysis (PCA) (Whibley *et al*. 2006) to identify and quantify the major axes of variation in abdominal expression patterns. Replicate images of pupal expression in each sex from each of the 35 reporter genes described above plus a negative control strain were used for this analysis. For each of these images, we marked the location of 30 pre-defined landmarks (blue dots in Figure 6A), used these landmarks to align the images, and extracted comparable fluorescence information from each of the images. A PCA was then run on this fluorescence data, and we analyzed the first three principal components (Supplementary Table 1), which together explained 55% of the variation.

The first principal component (PC1) explained 34% of the variation in expression among lines and primarily captured the overall fluorescence intensity (Figure 6A). Variability in this trait likely reflects differences in the average expression level driven by different fragments as well as some day-to-day variation in confocal conditions and the difference in ploidy for the five heterozygous lines. The second principal component (PC2), an orthogonal axis to PC1, explained 12% of the variance and primarily captured the intensity of fluorescence in abdominal segments A5 and A6 relative to A2-A4 (Figure 6A). Consistent with this interpretation, fragments classified as driving sexually dimorphic expression tended to have greater values of PC2 in males than females relative to fragments driving expression patterns not described as sexually dimorphic (Figure 6B, one-sided t-test, P = 0.0009). We note two exceptions to this relationship. First, the mel_A3 fragment drove high expression in abdominal segments A5 and A6 classified as sexually monomorphic (Figure 3A; Supplementary Figure 2) but PC2 was greater in males than females suggesting this fragment drives subtle differences between the sexes. Second, expression driven by the mel_A6 fragment was scored as sexually dimorphic (Figure 3A; Supplementary Figure 2) but PC2 was more similar in males and females than in other cases scored as dimorphic (Figure 3A; Supplementary Figure 2). In addition to sexual dimorphism in A5 and A6, PC2 also captured the prominence of stripes along the posterior edge of abdominal segments A2-A4, presumably because most fragments driving elevated expression in A5 and A6 also drove broad expression in A2-A4 (Figures 3, 4, 5). The third principal component (PC3) explained 9% of the variance and seemed to capture whether expression was elevated at the posterior edge of each segment (Figure 6A). The independent variation of these three traits (overall expression level, expression in A5 and A6, and abdominal stripes) suggests that they are controlled by distinct developmental processes.

**Figure 6.**
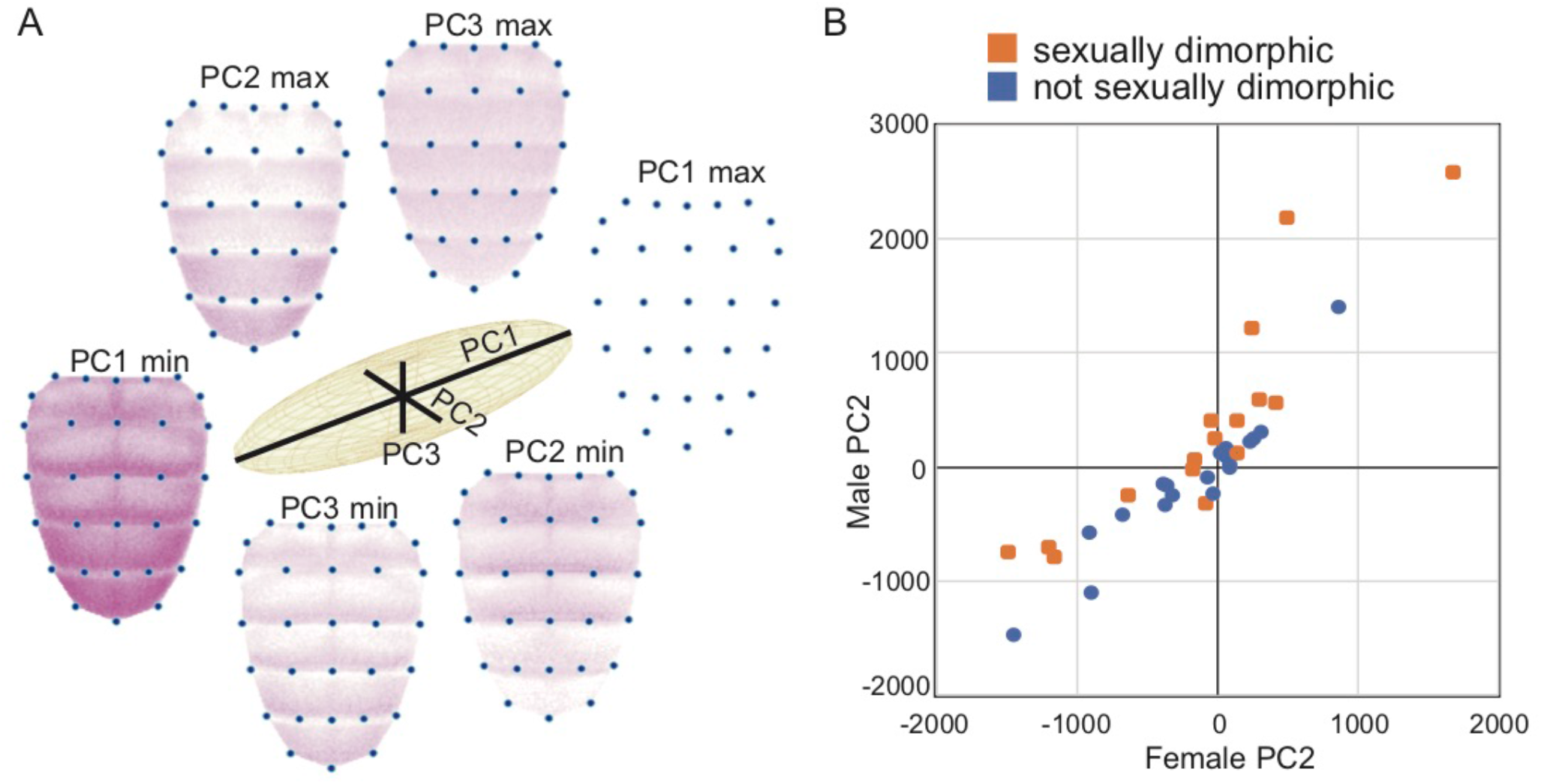
Principal components analysis of abdominal expression patterns. (A) Relative proportions of variation in abdominal expression patterns explained by principal components 1, 2, and 3 (PC1, PC2, PC3) are shown with black lines in the center of the figure. Abdominal images at the ends of each of these black lines show inferred expression patterns that correspond to the minimum (min) and maximum (max) values of each of the principal components. Comparing the images with min and max values for each PC shows the elements of abdominal expression captured by that principal component. (B) The scatterplot compares values of PC2, which primarily captures elevated expression in abdominal segments A5 and A6, between males and females for each reporter gene. PC2 values for reporter genes driving expression in abdominal segments A5 and A6 interpreted as sexually dimorphic are shown in orange and PC2 values for reporter genes driving sexually monomorphic expression in abdominal segments A5 and A6 are shown in blue.

### Conservation and divergence of enhancer activities are not well-reflected in conservation and divergence of *cis*-regulatory sequences

To search for DNA sequences driving specific elements of the expression patterns shown in Figures 3, 4, and 5, we examined sequence similarity among the 5’ intergenic and intronic regions from *D. melanogaster*, *D. pseudoobscura*, and *D. willistoni* (Figure 7). The program *promoterwise*, which was designed to search for sequences regulating transcription in orthologous non-coding sequences (Ettwiller *et al*. 2005), was used for this analysis. This program uses a local alignment method to identify regions of sequence with greater similarity than expected by chance and is robust to inversions and translocations as well as insertions and deletions. Sequences found to be more similar between species than expected by chance (Supplementary File 2) are connected with black lines in Figure 7.

**Figure 7.**
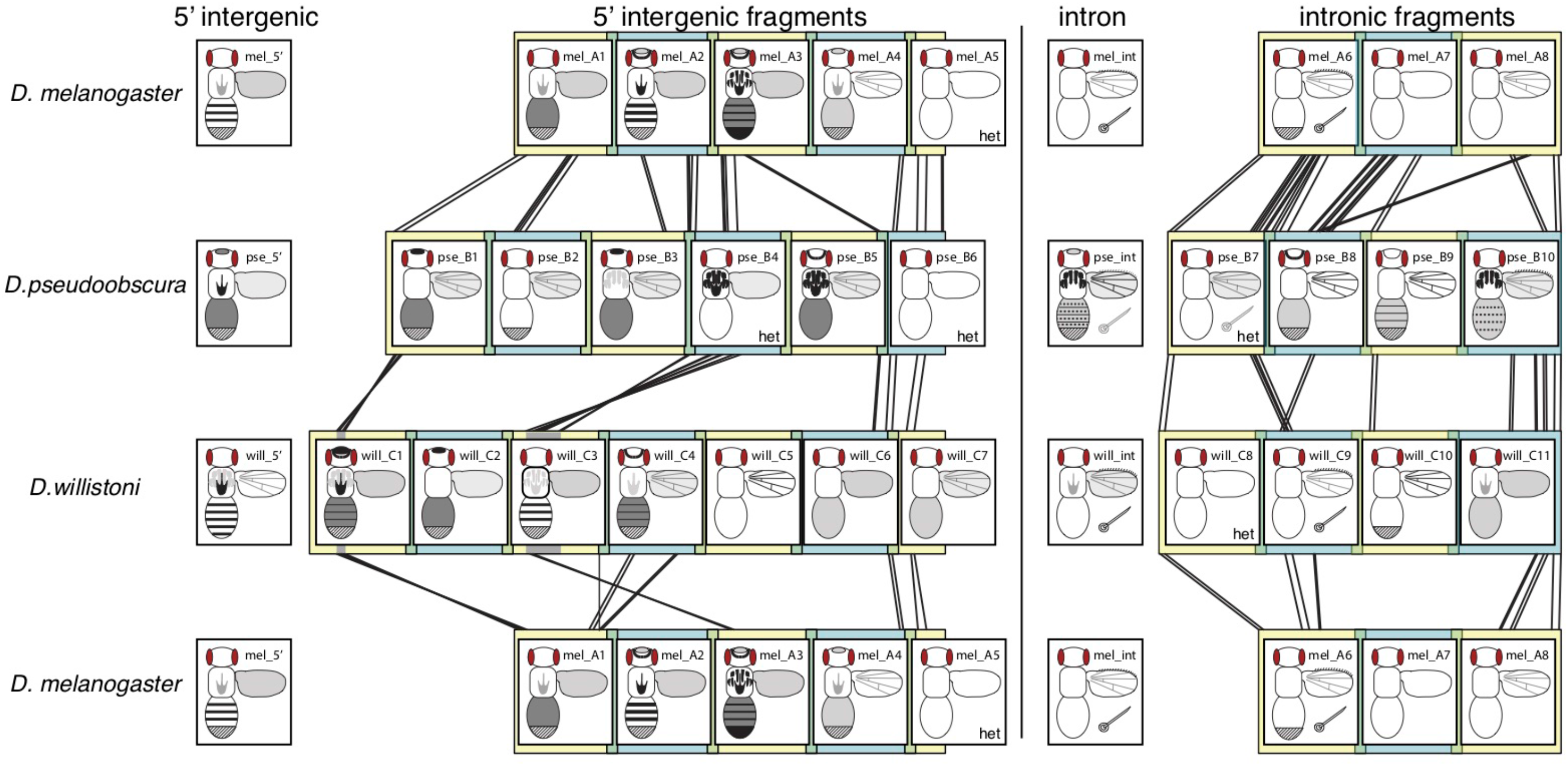
Summary of enhancer activity and sequence similarity for *yellow* fragments. Overlapping fragments from the 5’ intergenic and intronic regions of *D. melanogaster*, *D. pseudoobscura*, and *D. willistoni yellow* are show in alternating yellow and blue square blocks, with regions of overlap shown in green. Black lines between fragments from different species show regions of significant sequence similarity detected with the *promoterwise* program, as described in the Methods section. Enhancer activity observed from each of these fragments as well as the full 5’ intergenic and intronic sequences from each species are represented in schematics similar to that shown in Figure 2. Regions of the body shaded dark grey or black showed higher expression levels than regions of the body shaded light grey. The striped region at the posterior end of the abdomen indicates sexually dimorphic expression with elevated expression in abdominal segments A5 and A6 of males. The posterior region is fully darkened in the schematic for fragment mel_A3 because it drove elevated expression in abdominal segments A5 and A6 of both males and females. The five fragments assayed only in flies heterozygous for the reporter gene are marked with “het”.

As expected from the phylogenetic relationships among these three species (Figure 1A), the greatest sequence similarity was observed between *D. melanogaster* and *D. pseudoobscura*. We found no evidence of large translocations or rearrangements between any pair of species, consistent with (Kalay and Wittkopp 2010); the orthologous 5’ intergenic and intronic sequences appeared to be syntenic (Figure 7). Despite this overall synteny, we observed what appear to be two small inversions (~20bp & ~300bp) specific to the *D. willistoni* lineage (grey regions in will_C1 and will_C3 fragments, Figure 7). Sequence similarity was detected between all pairs of species near the 5’ and 3’ ends of both the 5’ intergenic and intronic fragments, consistent with the regions assayed being orthologous.

Comparing regions of sequence similarity between species to enhancer activities, we observed two cases where similar enhancer activities were driven by fragments with shared sequences. First, mel_A3, pse_B4, and will_C3, all drove expression in the flight muscle attachment sites and contained regions of sequence similarity suggesting a conserved enhancer inherited from the common ancestor shared by all three species. The pse_B5 fragment also drove expression in the flight muscle attachment sites and showed sequence similarity with mel_A3, but not will_C3. Expression in this region of the thorax was driven by pse_B10 and will_C1 as well, but neither of these fragments showed sequence similarity with any other fragment driving expression in the flight muscle attachment sites, suggesting that this similar enhancer activity may have evolved independently.

Second, in the intron, the two fragments that drove high levels of expression in bristles on both the body and wing (mel_A6 and will_C9) showed sequence similarity, suggesting the presence of a conserved bristle enhancer inherited from a common ancestor. These two fragments also showed sequence similarity with the pse_B7 fragment, but we did not score pse_B7 as driving bristle expression in the body or wing (Figure 4C). We hypothesized that bristle expression driven by pse_B7 might have been missed because of heterozygosity of the pse_B7 reporter gene. Using digital image enhancements, we did indeed find evidence of bristle expression in the body driven by pse_B7 (Supplementary Figure 5), and thus conclude that mel_A6, pse_B7, and will_C9 contain an orthologous bristle enhancer. Even with digital enhancements, bristle expression along the wing margin was detected only with the pse_B10 fragment (Supplementary Figure 5), which did not show any sequence similarity to mel_A6 or will_C9, suggesting that this activity evolved independently.

Enhancers from *D. melanogaster* and *D. willistoni* driving expression of abdominal stripes might also have a common origin, but this is less obvious from the sequence comparisons. The mel_A2 and will_C3 fragments drove the clearest abdominal stripes but shared no identified regions of sequence similarity. However, sequences similar to regions of will_C3 as well as the neighboring will_C4, which also drove expression in stripes, mapped to the mel_A1 and mel_A3 fragments flanking mel_A2, suggesting enhancer activities in this larger region driving abdominal stripes might be homologous. Sequence similarity of mel_A2 and mel_A3 to pse_B3 and pse_B4, as well as similarity of pse_B4 to will_C3, further supports the orthology of these stripe enhancers. In addition, although the full *D. pseudoobscura* 5’ intergenic sequence did not drive elevated expression in abdominal stripes, there was some evidence of elevated expression driven by the pse_B3 fragment in females (Figure 4). Enhancer activities driving elevated expression along the posterior edge of abdominal segments were also observed for fragments will_C1 and pse_B9, but these sequences shared no similarity with mel_A2, pse_B4, or will_C3 and thus appear to be independently evolved.

Expression in all of the other structures scored (i.e., top of head and head circumference, broad abdominal expression, sexually dimorphic A5/A6 expression, thoracic trident expression, and expression in the wing blade and veins) was driven by multiple fragments within each species without any clear patterns of sequence similarity linking these fragments (Figure 7). Of these traits, enhancers driving sexually dimorphic expression have been studied most extensively in prior work, with two transcription factors, Abdominal-B (Abd-B) and Bric-a-brac (Bab) shown to directly regulate a sexually dimorphic enhancer of *yellow* in *D. melanogaster* (Jeong *et al*. 2006; Camino *et al*. 2015; Roeske *et al*. 2018). We tested for enrichment of motifs representing binding sites for Abd-B, Bab, or any of ~400 other transcription factors within the 13 fragments that drove sexually dimorphic expression in A5 and A6 relative to the 16 fragments that did not. None of the 1419 motifs tested were enriched among the fragments that drove sexually dimorphic expression. We also failed to find significant enrichment of any of these motifs among fragments containing the putatively orthologous bristle or flight muscle attachment site enhancers described above.

Finally, we compared results from this study to data from a prior study using a yeast-one-hybrid assay to test 670 transcription factors for evidence of binding to these same *yellow* 5’ intergenic and intronic fragments (Kalay *et al*. 2016). Only one transcription factor, Hr78, showed evidence of binding to fragments from all three species in this prior work (mel_A2, mel_A3, mel_A4, mel_A5, pse_B1, and will_C1). The pse_B1 and will_C1 showed some sequence similarity, but none was detected between these fragments and any of the four fragments from *D. melanogaster* (Figure 7). Furthermore, we did not observe any expression patterns that were unique to this set of fragments (Figure 7).

## Discussion

By systematically testing enhancer activity of overlapping fragments in the 5’ intergenic and intronic regions of *yellow* from three Drosophila species, we have (1) identified modular enhancers within larger *cis*-regulatory regions, (2) observed many redundant enhancers with overlapping function, and (3) uncovered cryptic enhancer activities in all 5’ intergenic and intronic regions tested except the *D. pseudoobscura* intron. Each of these findings is discussed in more detail below. Taken together, these data show that regulation of the Drosophila *yellow* gene is more complex than suggested by prior work, and its *cis*-regulatory sequences have properties that might have facilitated the frequent changes in expression observed among species for this gene.

### *yellow* enhancer modules refined and redefined

Most previous studies of *Drosophila yellow cis*-regulatory sequences have focused on tissue-specific enhancer modules (reviewed in Massey and Wittkopp 2016; Rebeiz and Williams 2017), which are continuous regions of *cis-*regulatory DNA that drive a particular subset of a gene’s expression pattern. For *D. melanogaster yellow*, the locations of such modules were first suggested by deletion studies showing that specific regions of 5’ intergenic and intronic sequence were necessary for pigmentation of specific body parts (Geyer and Corces 1987; Martin *et al*. 1989). Wittkopp *et al*. (2002) later demonstrated that these regions identified as necessary for expression in the body and wing were also sufficient to drive expression in these tissues, consistent with the idea that *yellow* was regulated by a collection of modular, tissue-specific enhancers.

Early studies defined the “body enhancer” as a region containing most of the sequence from the mel_A3 fragment and the 5’ half of sequence from the mel_A4 fragment (Wittkopp *et al*. 2002, Supplementary Figure 6). However, we found that sequences in the mel_A2 fragment provided the best reproduction of body expression driven by the full 5’ intergenic sequence and seen for the native Yellow protein (Hinaux *et al*. 2018). The mel_A3 sequence drove broader and stronger expression than the full 5’ intergenic sequence throughout the abdomen, thorax, and head, and the mel_A4 sequence drove lower levels of broad expression without pronounced abdominal stripes (Figure 3, Supplementary Figure 2). These data are consistent with (Jeong *et al*. 2006), who found that extending the originally described “body enhancer” to include mel_A2 sequences and additional mel_A4 sequence (Supplementary Figure 6) produced a more faithful representation of *D. melanogaster yellow* expression in the abdomen.

The prominent abdominal stripes of *D. melanogaster yellow* expression were driven by both mel_A2 and mel_A3 (Figure 3), but because these two fragments overlap by 100bp, this expression pattern might be controlled by a single, contiguous, modular abdominal stripe enhancer. Prominent abdominal stripes of expression were also driven by orthologous sequences in the *D. willistoni* 5’ intergenic region, suggesting that this enhancer activity is conserved between these two species. Despite being more closely related to *D. melanogaster* than *D. willistoni*, orthologous sequences from *D. pseudoobscura* did not drive expression in abdominal stripes other than a faint pattern limited to females driven by the pse_B3 females. Rather, they drove expression broadly throughout the abdomen, which suggests that this sequence retained enhancer activity in the abdomen but evolved a new expression pattern within this tissue. Sequences in these orthologous regions (mel_A3; pse_B3, pse_B4, pse_B5; will_C3) also appear to have enhancer activity driving expression in the flight muscle attachment sites and throughout the wing blade that has been conserved in all three species (Figure 7). This observation is consistent with prior studies of the *yellow* “wing enhancer” in *D. melanogaster*, which defined the wing enhancer as a region of sequence including most of the mel_A2 fragment with some sequence from the 5’ end of mel_A3 (Wittkopp *et al*. 2002; Gompel *et al*. 2005) (Supplementary Figure 6). It is also consistent with the observation in Kalay and Wittkopp (2010) that enhancers controlling expression in the wing blade and body tend to be located in the same genomic region.

A modular enhancer driving sexually dimorphic expression that is higher in the A5 and A6 abdominal segments of males relative to females has also been reported for *D. melanogaster*. This activity was localized to a region of *yellow* 5’ intergenic sequence overlapping both mel_A3 and mel_A4 in two prior studies (Jeong *et al*. 2006; Camino *et al*. 2015), with binding sites for Abd-B (Jeong *et al*. 2006) and Bab-1 (Roeske *et al*. 2018) identified in and near the region of overlap between mel_A3 and mel_A4 (Supplementary Figure 6). Consistent with these data, we observed sexually dimorphic expression driven by the mel_A4 fragment as well as elevated expression in abdominal segments A5 and A6 driven by the mel_A3 fragment, which also showed signs of being sexually dimorphic in the PCA analysis (Figure 3, Supplementary Table 1). Fragments from the orthologous region of *D. pseudoobscura* did not drive sexually dimorphic expression, but those from *D. willistoni* did (Figure 7). This sexually dimorphic expression driven by fragments from *D. willistoni* changes our understanding of when the sexually dimorphic enhancer of *yellow* most likely evolved, as discussed further in the cryptic enhancer section below.

Finally, we identified a modular enhancer controlling expression in bristles. In all three species, bristle expression was driven by the *yellow* intron, with orthologous fragments from *D. melanogaster, D. pseudoobscura*, and *D. willistoni* (mel_A6, pse_B7 and will_C9) all driving expression in bristles on the body. These *D. melanogaster* and *D. willistoni* fragments (mel_A6 and will_C9) also drove expression in bristles on the wing. Surprisingly, no wing margin bristle expression appeared to be driven by pse_B7; rather, expression in wing bristles was driven by a non-orthologous fragment from the *D. pseudoobscura* intron, pse_B10. The co-localization of enhancers driving bristle expression in the body and wings of both *D. melanogaster* and *D. willistoni* suggests a common set of transcription factor binding sites might be activating expression in bristles of both tissues; however, the data from *D. pseudoobscura* suggests that it might also be possible to independently regulate expression in body and wing bristles.

### Redundant enhancers are common for the *yellow* gene

Although studies of enhancers have focused primarily on tissue-specific modules (Arnone and Davidson 1997; Carroll 2008; Rebeiz *et al*. 2009; Lorberbaum *et al*. 2016), it has become clear in the last decade that multiple enhancers driving overlapping expression patterns, known as redundant or “shadow” enhancers (Hong *et al*. 2008; Hobert 2010; Barolo 2012), are also common (Zeitlinger *et al*. 2007; Lagha *et al*. 2012; Miller *et al*. 2014; Cannavò *et al*. 2016; Osterwalder *et al*. 2018). Such redundant enhancers can be evolutionarily conserved because they confer robustness in the face of genetic variation or environmental variation, or have different functions in different tissues (Frankel *et al*. 2010; Perry *et al*. 2010; Fujioka and Jaynes 2012; Lam *et al*. 2015; Osterwalder *et al*. 2018). Redundant enhancers might also play an important role during evolution because they provide opportunities for evolutionary novelty (Hong *et al*. 2008; Perry *et al*. 2009; Levine 2010; Long *et al*. 2016).

We found that redundant enhancers were common for the *yellow* gene: expression in all structures scored except bristles, was driven by at least two non-overlapping fragments in at least one of the three species (Figure 7). For example, broad expression in the abdomen was driven by three fragments from *D. melanogaster*, six fragments from *D. pseudoobscura* and six fragments from *D. willistoni*. Redundant enhancers driving expression throughout the wing blade, in abdominal stripes, and in segments A5 and A6 of *D. melanogaster* males were especially surprising given that these patterns were previously described as being controlled by a single enhancer module (Wittkopp *et al*. 2002; Gompel *et al*. 2005; Prud’homme *et al*. 2006; Jeong *et al*. 2006; Camino *et al*. 2015; Roeske *et al*. 2018). In the wing blade, expression was driven by three, seven, and seven fragments from *D. melanogaster, D*. *pseudoobscura*, and *D. willistoni*, respectively (Figure 7). In *D. willistoni*, abdominal stripes of expression were driven not only by fragments orthologous to the *D. melanogaster* stripe enhancer, but also by the will_C1 fragment that did not show any sequence similarity to other fragments driving expression in abdominal stripes (Figure 7). Finally, in *D. melanogaster*, sexually dimorphic expression was driven not only by the previously characterized region overlapping the mel_A3 and mel_A4 fragments (Jeong *et al*. 2006; Camino *et al*. 2015; Roeske *et al*. 2018), but also by sequences in the mel_A1 fragment. Taken together, these data suggest that redundant enhancers controlling *yellow* expression are much more common than suggested by prior work. Such redundancy of enhancer activity might make mutations capable of altering activity of specific enhancers insufficient to cause detectable changes in pigmentation.

### Cryptic enhancer activities are also common for *Drosophila yellow*

In some cases, we observed expression patterns driven by fragments from within the 5’ intergenic or intronic region that were not driven by the full fragment from the corresponding region. This ectopic expression might be caused by genomic sequence surrounding the transgene insertion site, but a negative control reporter gene lacking any enhancer fragment failed to drive expression in these patterns (Figure 2B,C). Ectopic expression could also be caused by new transcription factor binding sites created at junctions between *yellow* fragments and the reporter gene; however, the similarity of ectopic expression patterns among reporter genes and to expression patterns driven by the full 5’ intergenic and intronic regions of *yellow* from other species suggests they are unlikely to be caused by randomly generated transcription factor binding sites. We therefore interpret this ectopic expression as being driven by cryptic enhancers, which are enhancer sequences that have the ability to drive expression in a particular pattern but are repressed by surrounding sequences in their native context. Such cryptic enhancers have been shown to be the source of novel expression patterns in fruit flies (Rebeiz *et al*. 2011) and humans (Prabhakar *et al*. 2008) and may facilitate the evolution of novel enhancer activities more generally.

We found that multiple cryptic enhancer activities were present in the *yellow* 5’ intergenic and intronic regions that drove expression in the abdomen, thorax, wings and/or head. For example, we did not observe expression in the wing veins driven by the full 5’ intergenic sequence from either *D. melanogaster* or *D. pseudoobscura* but did observe expression in the wing veins driven by multiple sub-fragments from these regions (Figures 3, 4, and 7). Similarly, the 5’ intergenic regions of *D. melanogaster* and *D. willistoni* drove expression in abdominal segments A2-A4 most prominently in stripes at the posterior edge of each segment, but multiple 5’ intergenic sub-fragments from both species drove expression much more broadly within these abdominal segments. These broad abdominal expression patterns were similar to the broad expression driven by the full 5’ intergenic sequence of *D. pseudoobscura*, suggesting that *D. pseudoobscura yellow* expression might have evolved by inactivating repressive elements inherited from the common ancestor shared with *D. melanogaster* and *D. willistoni*.

Perhaps the most interesting of these cryptic enhancers were those from *D. willistoni* driving sexually dimorphic expression in abdominal segments A5 and A6 of males similar to the sexually dimorphic pigmentation seen in *D. melanogaster*. Pigmentation is not sexually dimorphic in *D. willistoni* (Figure 1), and neither the full 5’ intergenic nor intronic regions of *D. willistoni yellow* drove expression in this pattern (Figure 5). Nonetheless, four fragments from the *D. willistoni* 5’ intergenic sequence and one fragment from the D. *willistoni* intronic sequence drove higher expression in segments A5 and/or A6 of males than females (Figure 5). This observation indicates that sequences capable of driving sexually dimorphic abdominal expression (at least in *D. melanogaster*) were already present in the common ancestor shared by all three species. The sexually dimorphic enhancer activity seems to therefore have evolved earlier than described in Jeong *et al*. (2006) and Camino *et al*. (2015), which concluded that it arose in the lineage shared by *D. melanogaster* and *D. pseudoobscura* after it diverged from the lineage leading to *D. willistoni*. This element was inferred to be present in *D. pseudoobscura* despite its monomorphic pigmentation because sequences from the 5’ intergenic region of *D. pseudoobscura*, as well as those from its close relative *D. subobscuara*, which also has monomorphic pigmentation (Jeong *et al*. 2006), drove elevated abdominal expression in males when transformed into *D. melanogaster* (Camino *et al*. 2015). Our data are consistent with these observations, as we found that elevated expression in male A5 and/or A6 segments was driven by two 5’ intergenic and two intronic fragments from *D. pseudoobscura* (Figure 4).

Although pigmentation of *D. pseudoobscura* is not dimorphic, we do not describe these sexually dimorphic enhancer activities in *D. pseudoobscura* as cryptic enhancers because similar patterns were driven by the full 5’ intergenic and intronic sequences from *D. pseudoobscura* (Figure 4). Similarly, we do not describe sequences in pse_B9 and pse_B10 driving stripes of expression at the anterior and posterior edges of each abdominal segment as cryptic enhancers even though they are not reflected in *D. pseudoobscura* pigmentation. This is because similar expression patterns were also driven by the full *D. pseudoobscura* intron (Figure 4). It remains to be seen whether these sexually dimorphic and abdominal stripe enhancers from *D. pseudoobscura* also drive expression in these patterns in their native species. If not, this observation would suggest that these enhancer activities are observed in transgenic *D. melanogaster* because of *trans*-regulatory divergence between *D. melanogaster* and *D. pseudoobscura* (Wittkopp *et al*. 2003).

### *cis*-regulatory architecture of *yellow* might facilitate expression divergence

Expression of the Drosophila *yellow* gene has become a model system for understanding how gene regulation evolves in part because it often shows differences among species that correlate with divergent pigmentation. Selection for divergent pigmentation is assumed to be driving this expression divergence, but the *cis*-regulatory architecture of *yellow* itself might make it especially amenable to expression divergence. Like many genes, *yellow* is controlled by multiple, tissue-specific enhancers that allow genetic changes to affect expression in one tissue without altering expression in others. But we also found that there are many redundant enhancer activities within *yellow cis*-regulatory sequences, increasing opportunities for expression to be modified. In addition, we found that many *yellow cis*-regulatory sequences have cryptic enhancer activities that are repressed by flanking sequences. Perhaps most interestingly, we found that cryptic enhancer activities from one species often drive expression in patterns similar to those seen in other species, suggesting that these cryptic enhancers might be latent enhancers with evolutionary potential. Additional work is needed to determine whether these putatively latent enhancer activities have indeed been the source of what appear to be novel expression patterns in other species.

## Materials and Methods

### Constructing reporter genes and transgenic lines

Putatively orthologous 5’ intergenic and intronic regions of *yellow* from *D. melanogaster*, *D. pseudoobscura* and *D. willistoni* described in Kalay and Wittkopp (2010) were subdivided using PCR into approximately 1000 bp fragments (except for mel_A5, pse_B6, will_C7, which were 423 bp, 641 bp, 345 bp long, respectively). Each fragment overlapped the flanking fragments by approximately 100 bp, although fragments at the 5’ and 3’ ends of a 5’ intergenic or intronic region overlapped only with the element that was following or preceding them, respectively within the region surveyed. PCR was conducted using a mix of Taq DNA polymerase and Phusion High-Fidelity DNA Polymerase (New England Biolabs, NEB) in order to prevent PCR-introduced mutations. Recognition sites for the Asc-1 (NEB) restriction enzyme were introduced to the ends of each PCR product using primers with 5’ Asc-1 tails (Supplementary Table 2). Subsequently, the PCR products for *yellow* enhancer fragments were sub-cloned into the sequencing vector pGEM-T (Promega), and sequenced using M13 Forward and M13 Reverse primers (See Supplementary File 1 for sequences). Sequence-confirmed *yellow* enhancer fragments were then sub-cloned into a piggyBac-attB vector (as described in Kalay and Wittkopp, 2010) using the Asc-1 unique site. Next, we cloned the coding sequence for a nuclear Enhanced Green Fluorescent Protein (nEGFP) downstream (3’) of each enhancer fragment using the Fse-1 (NEB) unique restriction site. Multiple attempts to construct this reporter gene for pse_B4 failed with these methods, thus we employed the GeneArt method (Thermo Fisher Scientific) to produce this construct. In all cases, the resulting construct was confirmed with diagnostic restriction digests, prepared in high concentration using Zyppy Plasmid Maxi kit, reconfirmed with diagnostic digest, and sent to either Genetic Services, Inc (Cambridge, MA) or Genetivision Corporation (Houston, TX) for injection into the attP-40 line of *D. melanogaster*. This is the same attP site used to insert the reporter genes containing the full 5’ intergenic and intronic regions from each species, as described in Kalay and Wittkopp (2010).

### Documenting Reporter Gene Expression

*D. melanogaster* lines homozygous for the reporter gene were constructed as described in Kalay and Wittkopp (2010). Briefly, for each transgenic line, transformant flies were crossed to a 2nd chromosome balancer line marked by the “curly wings” phenotype (Bloomington Stock Center #7197). Among the F_1_ progeny, flies with green fluorescing eyes from the Pax6-GFP transformation marker and curly wings were crossed to each other. Among the F_2_ progeny, flies that had green fluorescing eyes and flat wings were assumed to be homozygous for the transgene and were crossed to each other to create a stable homozygous line for the transgene. For five of the *yellow* enhancer fragments, mel_A5, pse_B4, pse_B6, pse_B7, and will_C8, multiple crosses failed to produce flies with straight wings (i.e., individuals homozygous for the transgene). Therefore, for those lines, we analyzed flies that were hemizygous for the transgene (i.e., green fluorescing eyes with curly wings). Pupal bodies and wings from each line were prepared for microscopy 70-80 hours after pupa formation (APF) as described in Kalay and Wittkopp (2010) and imaged immediately using Leica SP5 confocal microscope. Images of the same tissue (body or wing) were processed identically for all lines in Adobe Photoshop CS6. All fragments except mel_A5, mel_A7, pse_B6, and will_C8 showed expression in at least one of the structures scored under these conditions. However, only one of these fragments, mel_A7, was assayed in flies homozygous for the reporter gene; heterozygosity of reporter genes driven by the other three fragments might have caused us to underestimate their activity.

### Principal Component Analysis

We created an appearance model to evaluate the main trends of spatial variation in gene expression along the abdomen in five male and five female flies from each of the transgenic lines except pse_B4 and the full *D. willistoni* intron (will_int). For pse_B4, four images of females and three images of males were used, and for will_int, only a single image from each sex was used, because strains carrying these two reporter genes went extinct during the course of this project. In all cases, we analyzed downsized 72-ppi images in jpg format. We created a 30-point model template, using the AAMToolbox in Matlab environment (Whibley *et al*. 2006). The point model was designed to capture the abdominal segments A2 to A6. We then performed Procrustes superimposition by modifying rotation, translation, and size so that all of the images were fitted into the mean abdomen shape, encapsulating 6000 pixels. This procedure allowed us to compare the patterns of RGB pixel values after removing size variation and/or subtle variation in image orientation. Finally, the warped abdominal images were used to perform a Principal Component Analysis, extracting the main trends of variation in RGB values for each of the 6000 pixels, giving an appearance space, and calculating PC values for each individual fly (Supplementary Table 1), using the “Shape Model Toolbox Software”, as described in Whibley *et al*. (2006).

### Sequence analysis

The sequences of the *yellow* 5’ intergenic and intronic regions from *D. melanogaster*, *D. pseudoobscura*, and *D. willistoni* were compared using the promoterwise program from the wise2.4 package (Ettwiller *et al*. 2005). This program allows alignments of sequences that aren’t necessarily co-linear, allowing for inversions and translocations, in addition to insertions and deletions. A cut-off of at least 20 bits was used because bit scores of more than 20 bits are extremely rare when comparing random sequences (Ettwiller *et al*. 2005). A custom R script was used to mark regions with significant sequence similarity on the schematic of *yellow* sequences shown in Figure 7.

To test for enrichment of potential transcription factor binding site motifs among fragments with similar enhancer activity, we used the Analysis of Motif Enrichment (AME) tool (McLeay and Bailey 2010) with the “Combined Drosophila Databases” from the MEME suite (memesuite.org). In all, we tested 1419 motifs, ranging from 4 to 26 nucleotides long, that describe binding sites for ~60% of all *D. melanogaster* transcription factors (http://memesuite.org/db/motifs, July 2018), for enrichment among sequences driving similar expression patterns. For sexually dimorphic expression, the 13 fragments shown as driving sexually dimorphic expression in Figure 7 were compared to the 16 sub-fragments from the 5’ intergenic and intronic regions of *yellow* that did not. For bristle expression, sequences from the putatively orthologous mel_A6, pse_B7, and will_C9 fragments were compared to all other intronic fragments except pse_B10. pse_B10 was excluded from the control set because it drove expression in wing margin bristles. For the flight muscle attachment sites, the putatively orthologous mel_A3, pse_B4, and will_C3 fragments were compared to all other 5’ intergenic fragments except mel_B3, mel_B5, and will_C1. This three fragments were excluded from the control set because they also drove expression in this region of the thorax.

## Acknowledgements

We thank Jonathan Massey and Henry Ertl for comments on the manuscript. Funding for this work was provided by National Institutes of Health NRSA fellowship 5-F32-GM-119203 to JL, Programa de Apoyo a Proyectos de Investigación e Innovación Tecnológica grant from Universidad Nacional Autónoma de México (PAPIIT-UNAM IA200217) to UR, and grants from the National Science Foundation (MCB-1021398) and National Institutes of Health (R01 GM089736 and 1R35GM118073) to PJW.

---

**Supplementary Figure 1.**
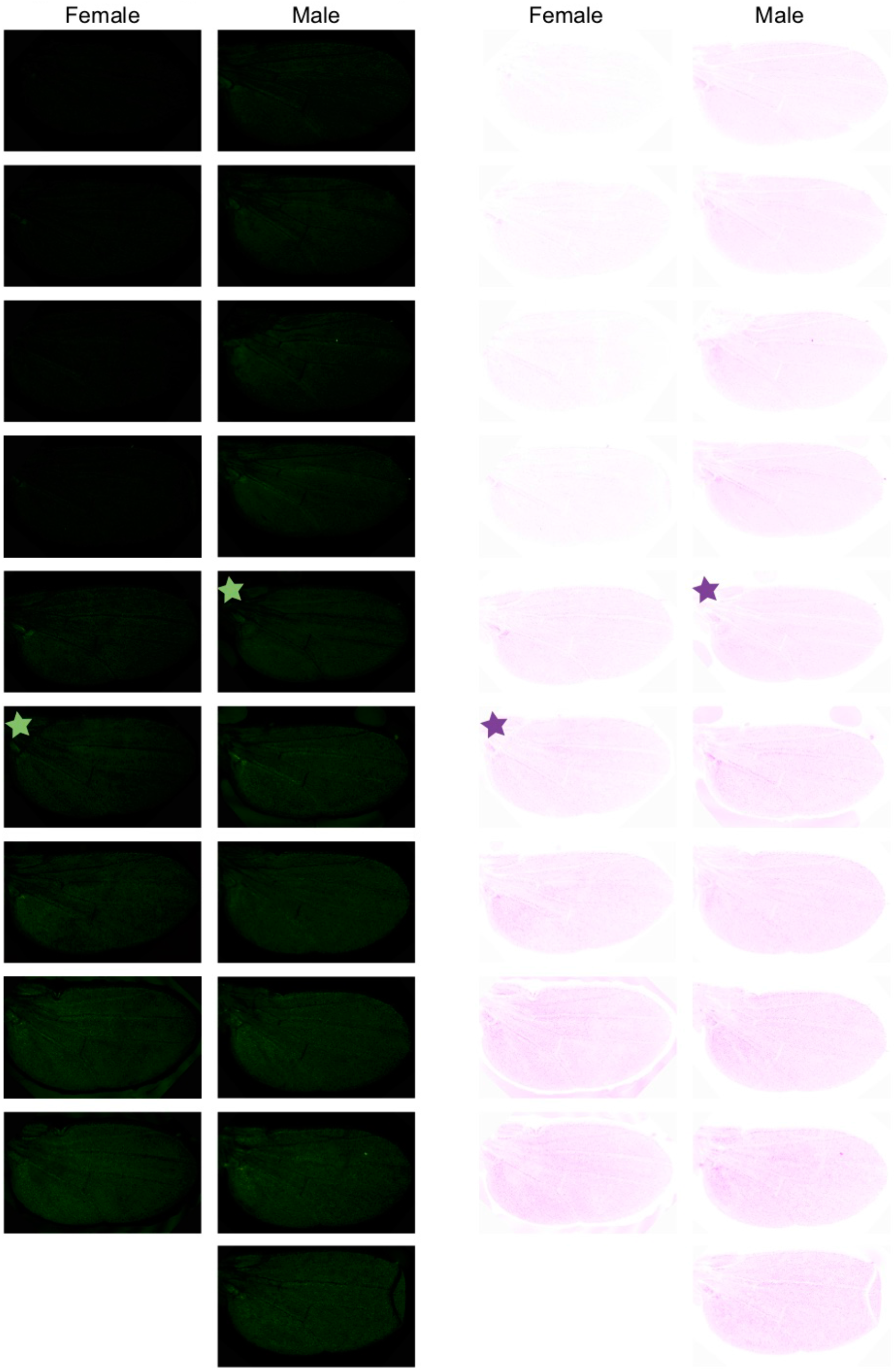
Variable levels of background fluorescence in wing blade. Fluorescence observed in a control strain carrying a reporter gene with only the basal promoter and GFP coding sequence is shown in the wing blade for nine female flies and ten male flies. Images in the two columns on the right show fluorescence using an inverted bolor scheme. Stars indicate the images included in Figure 2.

**Supplementary Figure 2.**
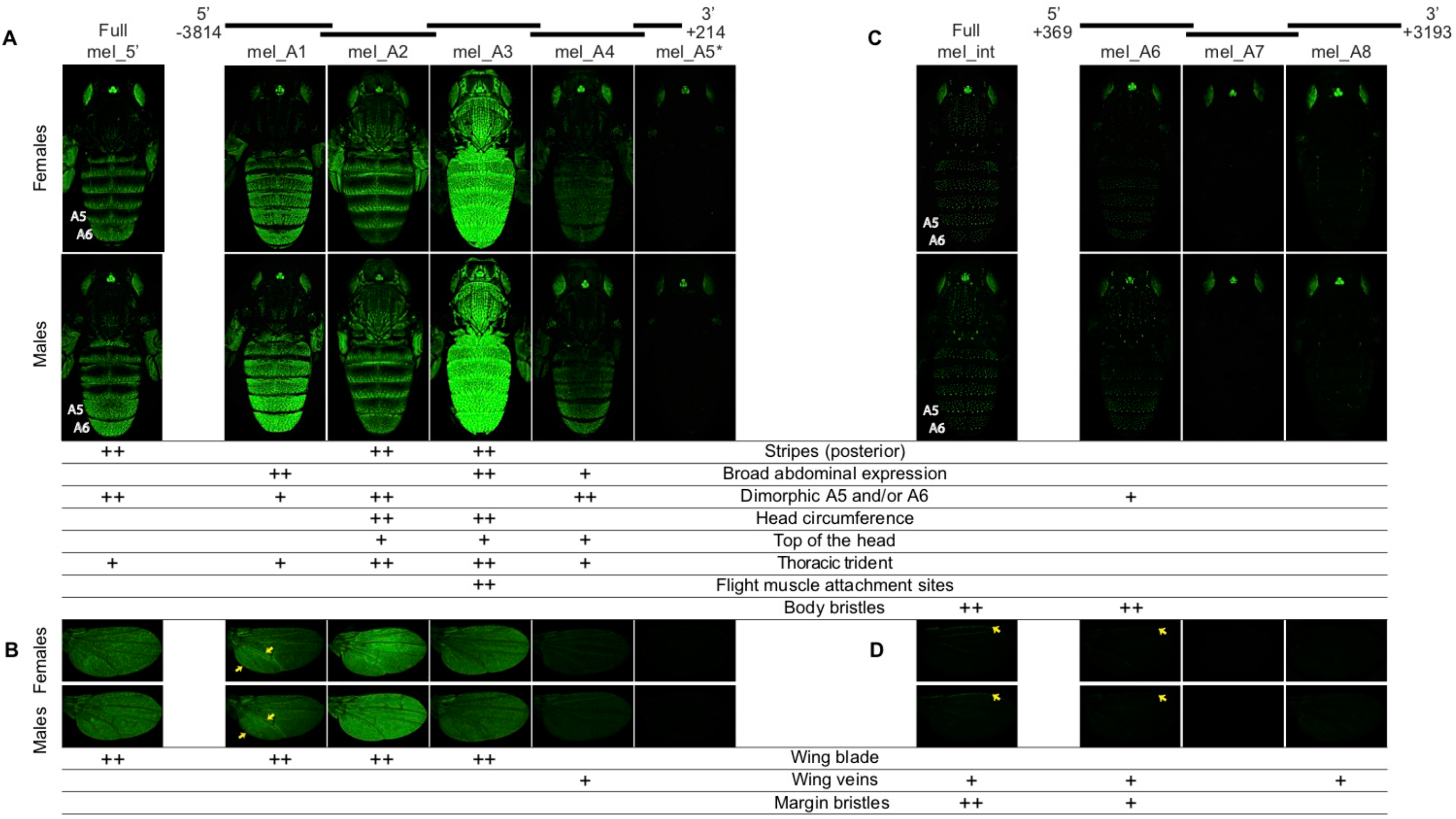
Enhancer activity of *D. melanogaster yellow* fragments. This figure shows the same information as Figure 3, but with an inverted color scheme that allows better visualization of high expression levels. All figure elements are as described in the Figure 3 legend except that the blue arrows in Figure 3 appear yellow here.

**Supplementary Figure 3.**
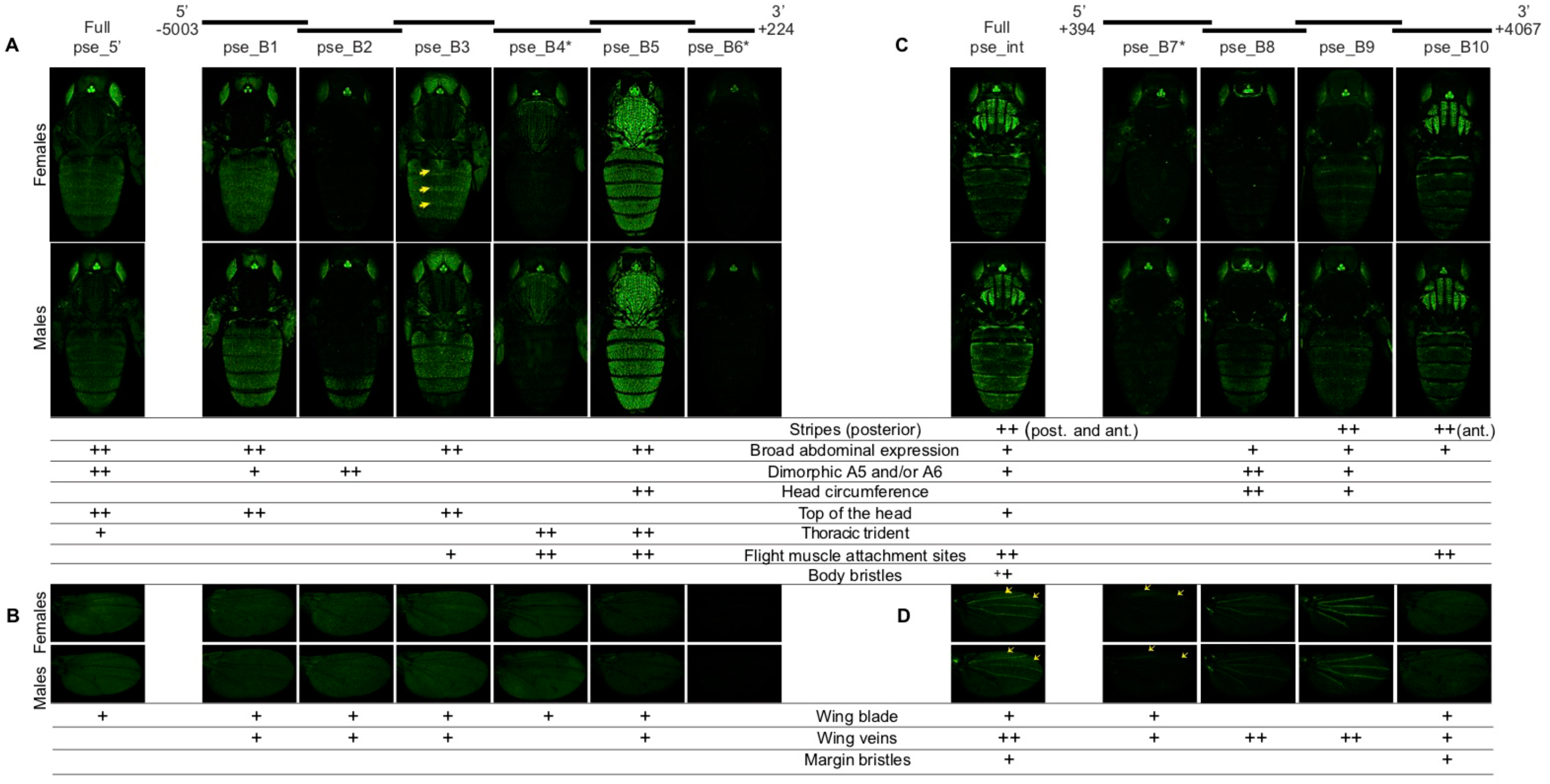
Enhancer activity of *D. pseudoobscura yellow* fragments. This figure shows the same information as Figure 4 but with an inverted color scheme that allows better visualization of high expression levels. All figure elements are as described in the Figure 4 legend except that the blue arrows in Figure 4 appear yellow here.

**Supplementary Figure 4.**
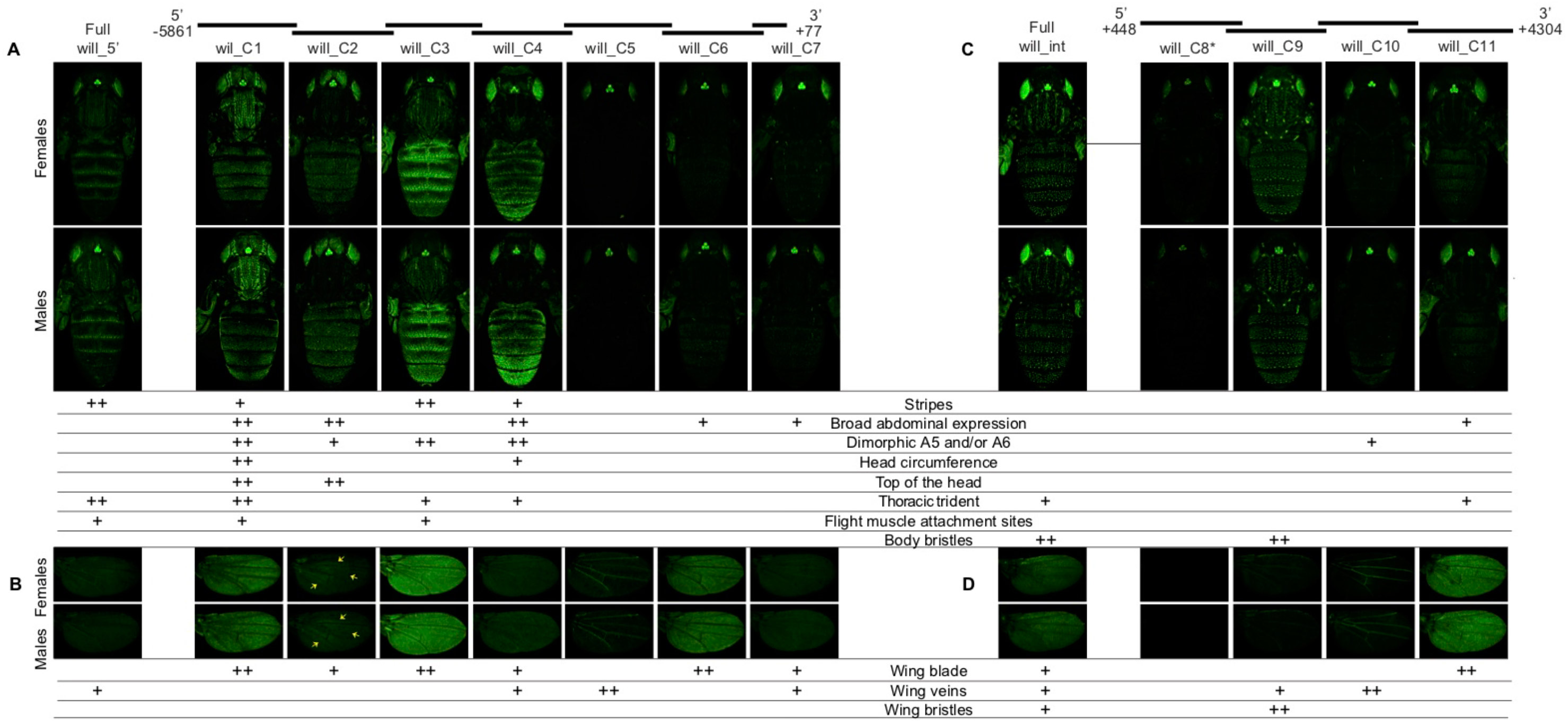
Enhancer activity of *D. willistoni yellow* fragments. This figure shows the same information as Figure 5 but with an inverted color scheme that allows better visualization of high expression levels. All figure elements are as described in the Figure 5 legend except that the blue arrows in Figure 5 appear yellow here.

**Supplementary Figure 5.**
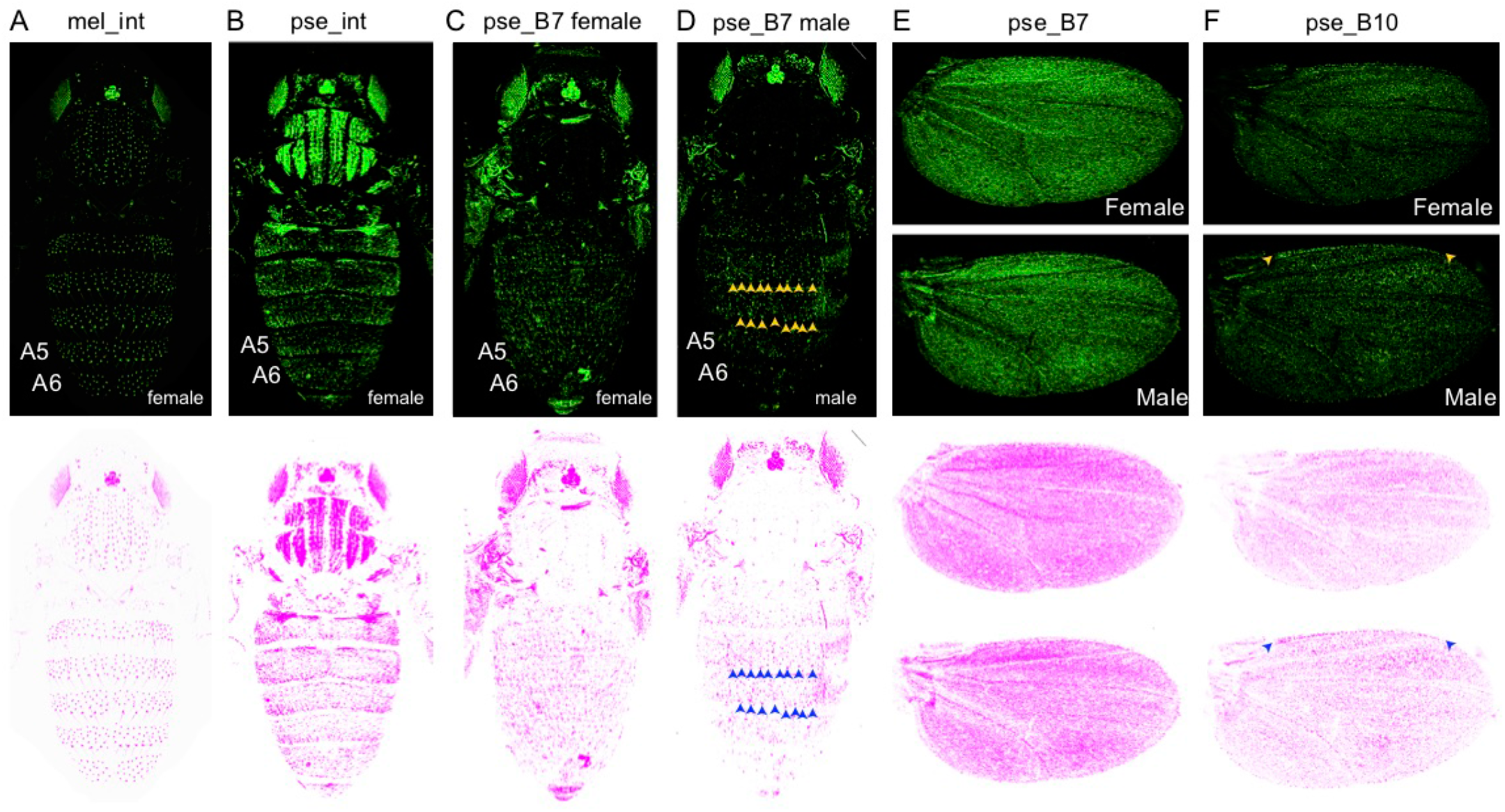
Image enhancement of fluorescence driven by fragment pse_B7. To search for enhancer activity in bristle cells driven by the pse_B7 fragment, which shares sequence similarity to fragments from *D. melanogaster* and *D. willistoni* that drive expression in body and wing bristles, we amplified signal in images of expression driven by the pse_B7 fragment in females (C) and males (D) in Photoshop. Expression driven by the *D. melanogaster intron* (A) and the full *D. pseudoobscura* intron (B) are shown for comparison in the body. Expression driven by the pse_B10 fragment is shown for comparison in the wing (E). The fluorescence signal reporting enhancer activity is shown in green in images in the top row and in magenta in images in the the bottom row. Arrowheads point to bristle expression in the body and wing margin of some fragments.

**Supplementary Figure 6.**
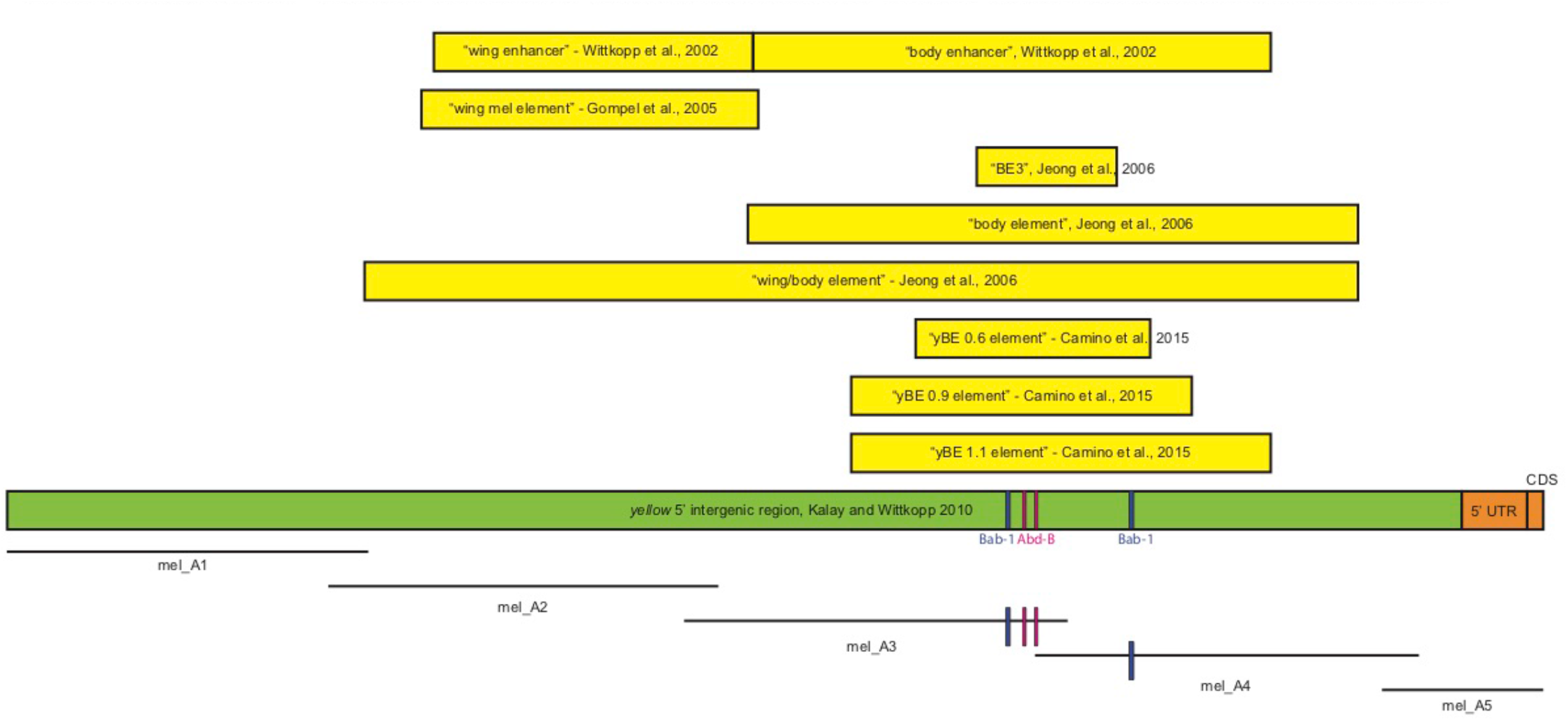
Summary of sequences from the *D. melanogaster yellow* 5’ intergenic sequence tested in prior work. Locations of experimentally confirmed binding sites for the Bric-a-brac (Bab-1) and Abdominal-B (Abd-B) transcription factors are indicated on the schematics of the full 5’ intergenic sequence as well as mel_A3 and mel_A4 fragments with blue and pink lines, respectively.

Supplementary Table 1 – Values of PC1, PC2, and PC3 for each of the reporter genes as well as negative control flies carrying a reporter gene without any enhancer sequences are shown.

Supplementary Table 2 – List of primers used to amplify each *yellow* 5’ intergenic or intronic sub-fragments from the three species in this study. Also included are, exact amplicon length and length of overlap with preceding and following fragments.

Supplementary File 1 – FASTA file with sequences of all *yellow* fragments tested for enhancer activity. Note that each sequence begins and ends with the recognition sequence for the AscI restriction enzyme (GGCGCGCC) used for cloning.

Supplementary File 2 – This .zip file contains *promoterwise* output and a custom R script used to process the *promoterwise* results from all pairwise comparisons.

